# deCLUTTER^2+^ pipeline to analyze calcium traces in a novel stem cell model for ventral midbrain patterned astrocytes

**DOI:** 10.1101/2022.11.08.515628

**Authors:** Martyna M. Grochowska, Federico Ferraro, Ana Carreras Mascaro, Domenico Natale, Amber Winkelaar, Valerie Boumeester, Guido J. Breedveld, Vincenzo Bonifati, Wim Mandemakers

## Abstract

Astrocytes are the most populous cell type of the human central nervous system and are essential for physiological brain function. Increasing evidence suggests multiple roles for astrocytes in Parkinson’s disease (PD), nudging a shift in the research focus, which historically pivoted around the ventral midbrain dopaminergic neurons (vmDANs). Studying human astrocytes and other cell types in vivo remains technically and ethically challenging. However, in vitro reprogrammed human stem cell-based models provide a promising alternative. Here, we describe a novel protocol for astrocyte differentiation from human stem cell-derived vmDANs-generating progenitors. This protocol simulates the regionalization, gliogenic switch, radial migration, and final differentiation that occur in the developing human brain. We have characterized the morphological, molecular, and functional features of these ventral midbrain astrocytes with a broad palette of techniques. In addition, we have developed a new pipeline for calcium imaging data analysis called deCLUTTER^2+^ (<u>dec</u>onvolution of <u>C</u>a<u>^2+^</u> f<u>lu</u>orescent pa<u>tter</u>ns) that can be used to discover spontaneous or cue-dependent patterns of Ca^2+^ transients. Altogether, our protocol enables the characterization of the functional properties of human ventral midbrain astrocytes under physiological conditions and in PD.

## 1. Introduction

Astrocytes are specialized glial cells essential for the healthy functioning of the nervous system (Sofroniew and Vinters, 2010). Astrocytes release neurotrophic factors (Dezonne et al., 2013; Gomes et al., 1999; Gomes et al., 2005; Nones et al., 2012), ensure the formation and maintenance of the blood-brain barrier (Verkhratsky and Nedergaard, 2018), regulate brain energy metabolism (Beard et al., 2021; Gonzalez-Reyes et al., 2017), modulate synaptic activity (Chung et al., 2015), and allow the movement of fluid between the paravascular spaces and the interstitium (Jessen et al., 2015). In addition, astrocytes provide metabolic support to neurons, including the uptake and exchange of mitochondria (Scheibye-Knudsen et al., 2015) and lipids (loannou et al., 2019; Liu et al., 2015).

The accumulating evidence points out that some of these homeostatic and neuronal health-promoting functions of astrocytes are impaired in Parkinson’s disease (PD) (Booth et al., 2017; MacMahon Copas et al., 2021). The majority of PD research focusses on the vulnerable ventral midbrain dopaminergic neurons (vmDANs), whose progressive loss is a disease hallmark. In recent years, however, the role of astrocytes in the neurodegenerative processes gained more attention. Studies have shown that genes known to cause inherited forms of PD are highly expressed in astrocytes and play vital roles in astrocyte function (Bandopadhyay et al., 2004a; Bandopadhyay et al., 2004b; di Domenico et al., 2019; Grochowska et al., 2021; Kim et al., 2016; Kim et al., 2013; Qiao et al., 2016; Strokin et al., 2012). Moreover, in PD, astrocytes can acquire a neurotoxic phenotype, enhance neurodegeneration and thus form a target for therapeutic intervention (Liddelow et al., 2017; Miyazaki and Asanuma, 2020; Schonhoff et al., 2020; Yun et al., 2018).

Astrocytes are a highly heterogeneous cell type based on their morphology, transcriptome, and physiology (Batiuk et al., 2020; Bayraktar et al., 2020; Oberheim et al., 2009; Pestana et al., 2020; Siletti et al., 2022). They elicit heterogeneous responses to injury and functionally specialize to their surrounding tissue (Bugiani et al., 2022). Astrocytes residing in the ventral midbrain are physiologically distinct from, e.g., astrocytes in the cortex and hippocampus (Siletti et al., 2022; Xin et al., 2019). Additionally, ventral midbrain astrocytes alleviate neuronal α-synuclein pathology, which is one of the main pathological hallmarks of PD (Yang et al., 2022).

Studying specific characteristics of human ventral midbrain astrocytes in PD remains challenging due to limited access to primary human cells from the affected brain regions. However, using induced pluripotent stem cell (iPSC) technology, it is now possible to generate various astrocyte populations of human origin in vitro (Lanjewar and Sloan, 2021). Evidence supports that patterning of iPSC-derived progenitors to rostral-caudal and dorsal-ventral identities with the same morphogens used for the neuronal subtype specification generates region-specific astroglial subtypes (Krencik et al., 2011).

A limited number of protocols exists that allows the generation of ventral midbrain astrocytes (**Supplementary Table 1**). A feature shared by these protocols is the generation of progenitor cells patterned toward a mesencephalic fate (floor-plate progenitors) that can also differentiate into vmDANs. Although these models have been an advancement in the PD research field, modifications, and improved characterization remain necessary to generate more accurate models of human ventral midbrain astrocytes.

Here, we describe a novel protocol for generating ventral midbrain patterned astrocytes from iPSCs. Similar to previous studies, we rely on the ventral neural tube patterning of iPSC-derived neuroepithelial cells to generate ventral midbrain astrocytes from a population of progenitors that can also produce vmDANs (Reinhardt et al., 2013). Our protocol mimics gliogenesis in vitro, including regionalization, gliogenic switch, radial migration, and final differentiation. In comparison to previous studies, our simplified protocol efficiently generates mature ventral midbrain astrocytes relatively fast, without the use of serum or transgenic reporters, and a limited number of necessary additives. We characterized these astrocytes using immunocytochemistry (ICC), RNA-sequencing (RNA-seq), and a battery of functional assays, including pro-inflammatory cytokine treatment, glutamate uptake, and fluorescence time-lapse calcium confocal imaging. Furthermore, we developed a novel pipeline for the functional analysis of calcium imaging data from in vitro experiments that we named deCLUTTER^2+^ (<u>de</u>convolution of <u>C</u>a<u>^2+^</u> f<u>lu</u>orescent pa<u>tter</u>ns). Altogether, our model can potentially study the properties of ventral midbrain astrocytes under physiological conditions and in PD.

## 2. Materials and methods

### 2.1 Primary cell lines

We generated eight human iPSC lines (**Supplementary Table 4**). Clonal lines D1-c1-iPSC and D1-c2-iPSC were generated from the commercially available dermal fibroblasts from a female donor (Gibco^™^, Lot number: 1903939). The skin biopsy was sampled at the age of 33. Clonal lines D2-c1-iPSC and D2-c2-iPSC were derived from the commercially available dermal fibroblasts from a female donor (Gibco^™^, Lot number: 181388). The skin biopsy was sampled at the age of 34. Clonal lines D3-c1-iPSC, D3-c2-iPSC, D3-c3-iPSC were generated from the primary fibroblasts from a female donor available in our institute. The skin biopsy was sampled at the age of 68. Line D4-c1-iPSC was derived from the primary cultures of erythroid progenitors (Leberbauer et al., 2005; van den Akker et al., 2010), which were generated from the peripheral blood of a female donor available in our institute. The blood was sampled at the age of 83. All study procedures were approved by the medical ethical committee of Erasmus MC and conformed to the principles of the WMA Declaration of Helsinki and the Department of Health and Human Services Belmont report. Participating subjects provided written informed consent for the use of the material for research purposes.

### 2.2 Generation of iPSC lines

Reprogramming of all primary cell lines into iPSCs used in this study was performed by the Erasmus MC iPS Core Facility. Primary cell lines were reprogrammed using the CytoTune^™^-iPS 2.0 Sendai Reprogramming Kit (Thermo Fisher Scientific) based on a modified non-transmissible form of Sendai virus (SeV), which contains reprogramming Yamanaka factors, OCT3/4, SOX2, KLF4, and C-MYC. The emerging iPSC colonies were manually picked and expanded four to five weeks after transduction. The selection was based on morphology. iPSCs were cultured in StemFlex^™^ medium (Gibco^™^) on Geltrex^™^ (Thermo Fisher Scientific) coated plates at 37 °C/5% CO_2_. iPSCs were passaged when they reached ~80% confluence using Versene (Gibco^™^). Multiple clones per line were assessed for their karyotype, expression of endogenous pluripotency factors, and three lineages (ectoderm, endoderm, and mesoderm) differentiation potential following the procedures of the Erasmus MC iPS Core Facility.

### 2.3 Primary and secondary antibodies

The primary antibodies used in this study are listed in Supplementary Table 3. Alexa Fluor^®^ 488 donkey anti-goat/mouse/rabbit, Alexa Fluor^®^ 594 donkey anti-mouse/rabbit, Alexa Fluor^®^ 647 donkey anti-rabbit, and Alexa Fluor^®^ 647 goat anti-guinea pig (all from Jackson ImmunoResearch) have been used as secondary antibodies for ICC experiments and microscopic analysis.

### 2.4 Characterization of iPSC lines with immunocytochemistry (ICC)

iPSCs grown on coverslips were fixed with 4% PFA for 15 min at room temperature (RT). Cells were washed three times for 5 min with 1× PBS buffer. Next, cells were incubated with ice-cold methanol for 10 min and washed one time with 1× PBS buffer. Subsequently, cells were permeabilized with 0.1% Triton-X100 (Sigma-Aldrich) in 1× PBS buffer for 10 min. Cells were blocked in a blocking buffer containing 1% BSA (Sigma-Aldrich) in 1× PBS-0.05% TWEEN^®^ 20 (Sigma-Aldrich). Then, cells were incubated with the blocking buffer containing primary antibody mixtures overnight at 4 °C. The next day, cells were washed three times for 5 min in 1× PBS-0.05% TWEEN^®^ 20 buffer. Cells were then incubated with appropriate Alexa Fluor^®^ secondary antibodies (1:500 dilution) for 1 hour at RT. Cells were washed three times with 1× PBS-0.05% TWEEN^®^ 20 buffer. Finally, coverslips were mounted with ProLong Gold with DAPI (Invitrogen). All stainings were imaged with Leica SP5 AOBS confocal microscope. Each image was detected on the spectral PMT detector with an HCX PL APO CS 40×/1.25 or HCX PL APO CS 63×/1.4 lens. Sections were irradiated with the following lasers: 405 diode UV, argon laser, DPSS 561, and HeNe 633, depending on the fluorophore combination. Scanning was performed at 400 Hz with a pixel size of 0.12 μm in the x, y-direction and 0.35 μm in the z-direction. The pinhole size was set to one airy unit (AU).

### 2.5 Generation of neural progenitor cells

Neural progenitor cells were generated via inhibiting BMP and TGFβ signaling (dual-SMAD) and stimulation of WNT and SHH signaling with small molecules according to published protocols (Grochowska et al., 2021; Reinhardt et al., 2013) with few modifications. iPSCs grown in feeder-free conditions (StemFlex^™^ in combination with Geltrex^™^) were detached from the plates using the Versene solution. Cells were dissociated as clumps and plated on irradiated mouse embryonic fibroblasts (MEFs) in StemFlex^™^ medium supplemented with 1× RevitaCell^™^ in a splitting ratio of 1:10. Cells were cultured at 37 °C/5% CO_2_ until the iPSC colonies reached the appropriate size in StemFlex^™^ media. Unwanted differentiations that arose around the iPSC colonies were manually removed. When colonies populated ~70% of the culture dish, cells were detached as clumps from the MEFs using Versene. Next, pieces of colonies were resuspended in Human Embryonic Stem Cell medium [HESC, 80% DMEM/F-12, 20% KnockOut Serum Replacement, 1% L-glutamine, 1% Penicillin-Streptomycin, 1% MEM-NEAA (all from Gibco^™^), and 0.0007% 2-β-Mercaptoethanol (Sigma-Aldrich)] supplemented with 10 μM SB-431542 (SB, Tocris), 1 μM dorsomorphin (DM, Abcam), 3 μM CHIR99021 (CHIR, Sigma), and 0.5 μM Purmorphamine (PMA, Stem Cell Technologies). Clumps of cells were then transferred to 10 cm Petri dishes and cultured in suspension for six days on a shaker at 80 RPM at 37 °C/5% CO_2_. On day two, the medium was replaced by N2B27 medium [DMEM/F-12– Neurobasal in 1:1 ratio, 1:100 B27 w/o Vitamin A, 1:200 N2, and 1% Penicillin-Streptomycin (all from Gibco^™^)] supplemented with 10 μM SB, 1 μM DM, 3 μM CHIR, and 0.5 μM PMA. On day four, the medium was changed to N2B27 medium supplemented with 3 μM CHIR, 0.5 μM PMA, and 150 μM Ascorbic Acid (AA, Sigma). On day six, embryoid bodies showing a developing neuroepithelium were collected, dissociated into smaller pieces, and plated on Corning^®^ Matrigel^®^ coated 12-well plates in N2B27 medium supplemented with 3 μM CHIR, 200 μM AA, and 0.5 μM PMA at 37 °C/5% CO_2_. Cell splits were usually performed at 1:10 or 1:20 ratios. After passage five, 0.5 μM PMA was replaced by 0.5 μM Smoothened Agonist (SAG, Abcam). Progenitors were passaged at least five times before astrocyte differentiations. Progenitors can be expanded in bulk (up to passage 30) and frozen for long-term storage.

### 2.6 Differentiation of neural progenitors into ventral midbrain astrocytes

Neural progenitors that reached 70-80% confluence were dissociated into single cells using Accutase^®^ (Sigma-Aldrich). Cells were seeded in ultra-low-attachment 96-well U-bottom plates (BIOFLOAT^™^) at the concentration of 15,000 cells per well in the N2B27 medium supplemented with 3 μM CHIR, 0.5 μM SAG, and 200 μM AA. Before placing in the incubator, plates were spun down at 220 × g for 5 min. Cells were cultured at 37 °C/5% CO_2_. After three days, progenitor cells formed spheres of size around 300-400 μm. On day three, the medium was replaced with a glial expansion medium containing DMEM/F-12, GlutaMAX, 1:50 B27 w/o Vitamin A, 1:100 N2, 1× MEM non-essential amino acids solution (NEAA), 1% Penicillin-Streptomycin (all from Gibco^™^), freshly supplemented with 10 mM HEPES (Gibco^™^), 10 ng/mL EGF (Peprotech), and FGF-2 (Peprotech). The medium was refreshed every other day. Cells were kept at 37 °C/5% CO_2_. On day 17, the medium was switched to a glial induction medium containing DMEM/F-12, GlutaMAX, 1:50 B27 w/o Vitamin A, 1:100 N2, 1× MEM non-essential amino acids solution (NEAA), 1% Penicillin-Streptomycin (all from Gibco^™^), freshly supplemented with 10 mM HEPES (Gibco^™^), 10 ng/mL EGF (Peprotech), and 10 ng/mL LIF (Peprotech). The medium was refreshed every other day until day 31. Cells were kept at 37 °C/5% CO_2_. On day 31, 15-20 spheres per line were carefully taken up and plated onto one well of a 6-well plate coated with Corning^®^ Matrigel^®^. Generally, spheres easily attach to the coated surface. After attachment to the coated plates, the cells migrate out of the spheres. From day 31, the cultures were maintained in glial maturation media containing DMEM/F-12, GlutaMAX, 1:50 B27 w/o Vitamin A, 1:100 N2, 1× MEM non-essential amino acids solution (NEAA), 1% Penicillin-Streptomycin (all from Gibco^™^), freshly supplemented with 10 mM HEPES (Gibco^™^), and 10 ng/mL CNTF (Peprotech). The medium was refreshed every other day until day 59. From day 60 onwards, cells were refreshed every two days with a glial maintenance medium containing DMEM/F-12, GlutaMAX, 1:50 B27 w/o Vitamin A, 1:100 N2, 1× MEM non-essential amino acids solution (NEAA), 1% Penicillin-Streptomycin, freshly supplemented with 10 mM HEPES (all from Gibco^™^). From day 60, cells were used in the downstream functional experiments. Cells were plated onto new Matrigel^®^ coated wells at 50.000 cells/cm^2^ when they repopulated the entire well. Detailed cell culture media composition is listed in **Supplementary Table 2**.

### 2.7 Characterization of iPSC-derived progenitors and astrocytes with ICC

Cells were fixed with 4% PFA for 15 min at RT. Cells were washed three times for 5 min with 1× PBS buffer. Next, cells were incubated in a staining buffer [50 mM Tris.HCl (pH 7.4), 0.9% NaCl, 0.25% gelatine, H2O] containing primary antibody mixtures overnight at 4 °C. The next day, cells were washed three times for 5 min in 1× PBS-0.1% TWEEN^®^ 20 buffer. Cells were then incubated with appropriate Alexa Fluor^®^ secondary antibodies (1:200 dilution) for 1 hour at RT. Secondary antibodies were washed three times with 1× PBS-0.1% TWEEN^®^ 20 buffer. Finally, cells were mounted with ProLong Gold with DAPI (Invitrogen). Each image was detected on the spectral PMT detector with an HCX PL APO CS 40×/1.25 or HCX PL APO CS 63×/1.4 lens. Sections were irradiated with the following lasers: 405 diode UV, argon laser, DPSS 561, and HeNe 633, depending on the fluorophore combination. Scanning was performed at 400 Hz with a pixel size of 0.12 μm in the x, y-direction and 0.35 μm in the z-direction. The pinhole size was set to 1 airy unit (AU). All images were processed using Fiji (Fiji Is Just ImageJ) software (version 1.53c) (Schindelin et al., 2012). For the quantitative characterization of neural progenitors and astrocytes, fluorescence-based thresholding was applied for each marker. The cell was considered positive for a marker if its fluorescent signal was above that threshold and within the boundaries of its belonging cell.

### 2.8 RNA extraction, sequencing, and alignment to reference genome

RNA was extracted from 14 iPSC-derived lines (eight progenitors and six astrocytes) generated from four individuals (**Supplementary Table 4**). Cells were harvested and lysed with lysis buffer (1 mL RTL+10uL β-mercaptoethanol) and scraped with a polypropylene disposable cell scraper. Cell lysates were transferred to 1.5 mL tubes and vortexed to dissolve possible cell clumps. Samples were further processed using the QIAGEN RNeasy^®^ (GTIN 04053228006121, LOT 172019069, REF 74106) Mini Kit according to the manufacturer’s instructions. RNA fragments were analyzed using the RNA 6000 Nano Kit on an Agilent 2100 Bioanalyzer instrument (Agilent Technologies) to determine the RNA integrity number (RIN). About 1 μg high-quality RNA sample (average RIN 9.9; **Supplementary Table 4**) from each sample was further processed with the Illumina NEBNext Ultra II Directional RNA Library Prep Kit (NEB #E7760S/L) according to the manufacturer’s instructions. The generated libraries were sequenced on an Illumina NovaSeq6000 with 150 bp paired-end reads and an average of 60 million reads per sample (GenomeScan, Leiden, Netherlands). RNA-seq data quality was assessed using fastqc v0.11.9 (Andrews, 2010) and summarized using MultiQC v1.12 (Ewels et al., 2016). RNA-seq reads were aligned to the GRCh38 human reference genome with STAR v2.7.10 (Dobin et al., 2013) run in multisample 2-pass mapping mode to improve the detection of novel splice junctions. The counts of reads per gene were determined using FeatureCounts v2 (Liao et al., 2014) using ENSEMBL gene annotations v107.

### 2.9 Cell lines karyotyping

RNA-seq data were processed according to the GATK v4.2.5 guidelines (McKenna et al., 2010) to generate a VCF with HaplotypeCaller. The possible presence of chromosomal multiplications was then assessed using eSNP-Karyotyping (Weissbein et al., 2016) with the BAM and VCF in combination with dbSNP v155. As previously described, the allelic ratio in sliding windows of 151 SNPs with a depth minimal allele frequency above 0.2 and covered by more than 20 reads were compared with that of the rest of the genome. Significant multiplications were identified as those with a false discovery rate < 0.05.

### 2.10 RNA-seq expression data analyses

Digital expression matrices of the analyzed cell lines were analyzed in R v4.2.1. Established cell-type specific markers were obtained from BRETIGEA v.1.0.3 (McKenzie et al., 2018). Moreover, we used previously described markers to characterize precursor cells and the maturity of the astrocytes (Sloan et al., 2017). Low-expressed genes (<10 cumulative raw counts across all samples) were filtered out, resulting in 28,075 expressed genes. DESeq2 v1.36.0 (Love et al., 2014) was used to transform raw counts, assess sample-to-sample distances and clustering via principal component analysis (PCA) on rlog-transformed data, and perform differential expression analysis. The obtained *p*-values were corrected with the Benjamini-Hochberg (BH) method (*BHp*-value), and differentially expressed genes (DEGs) were identified according to the following thresholds: *BHp*-value< 0.05 and |fold change| > 1.5.

### 2.11 RNA-seq functional analysis

Gene set enrichment analysis (GSEA) was performed using the clusterProfiler v4.4.4 (Wu et al., 2021) and org.Hs.eg.db v3.15.0 R packages with the Gene Ontology (GO) database. GO terms were filtered to include sets comprised of 20 - 2,000 genes. The genes in our datasets were ranked in descending order by the negative logarithm in base 10 of the adjusted *BHp*-value, multiplied for the sign of the fold change. All resulting *p*-values were corrected using the BH method. A nervous system-centric analysis was conducted extracting the 181 terms surviving out filtering criteria out of the 1,256 child terms of the nervous system development set (GO:0007399).

### 2.12 Pro-inflammatory cytokine treatment: A1-like reactivity assay

We adapted the reactivity assay from Barbar et al (Barbar et al., 2020) to obtain an A1 reactive astrocyte phenotype. Astrocytes were plated onto new Matrigel^®^ coated wells at 50.000 cells/cm^2^. Two days later, astrocytes were treated with 30 ng/mL TNF-α (tumor necrosis factor α), 3 ng/mL IL-1α (interleukin 1α), and 400 ng/mL C1q (complement component 1, subcomponent q) for 24 hours. After 24 hours, cells were collected, and the total RNA was isolated using the RNeasy Mini Kit (Qiagen) as recommended by the manufacturer. For each sample, on-membrane DNase I (RNase-Free DNase Set, Qiagen) digestion was performed according to the manufacturer’s protocol. The integrity of the total RNA was assessed using agarose gel stained with GelRed^™^ (Biotium) and spectrophotometric analysis using NanoDrop 2000/2000c. Sharp, clear 28S and 18S rRNA bands without smearing and 260/280 values of ~2.0 indicated intact and pure RNA. Next, 0.5 μg of the total RNA was converted into cDNA using the iScript cDNA Synthesis Kit (Bio-Rad). qPCR was performed on CFX Opus 96 Real-Time PCR System (Bio-Rad) with ~0.1 μg cDNA per reaction. The following cycling conditions were used: 3 min at 95 °C (initial denaturation), 40 cycles of 5 s at 95 °C, and 30 s at 60 °C. Data analysis was performed using CFX Maestro software 2.3 (Bio-Rad). Briefly, the normalized expression of each target gene was calculated using the delta–delta Cq method (Livak and Schmittgen, 2001). Relative mRNA levels for *C3, LCN2, SERPINA3*, and *GFAP* were determined after normalization to the geometric mean of the following housekeeping genes: *COPS5, CLK2*, and *RNF10*. All assays, spanning at least one intron, were validated by demonstrating linearity over 3 orders of magnitude and by observation of a single melt peak by plotting RFU data collected during a melt curve as a function of temperature. Primers were adapted from previously published articles or designed using Primer3 v. 0.4.0 online tool (https://bioinfo.ut.ee/primer3-0.4.0/) and are listed in **Supplementary Table 5**.

### 2.13 Glutamate uptake assay

Astrocytes grown in 6-well plates were refreshed with glial maintenance media or glial maintenance media supplemented with 100 μM L-Glutamic Acid (Tocris) for 3 h. The glutamate concentration in astrocyte cultures was determined using the Glutamate Assay kit (Sigma-Aldrich, MAK004). To obtain cell homogenates, astrocyte cultures were lifted with Accutase^®^, briefly centrifuged, and resuspended with 100 μL of the Glutamate Assay Buffer. Samples were centrifuged at 13,000 × g for 10 min to remove insoluble material. Enzymatic reaction mixes were prepared by mixing Glutamate Assay Buffer, Developer, and Enzyme Mix according to the manufacturer’s specifications and were subsequently mixed with 50 μL of cell homogenates and incubated for 30 min at 37 °C. Absorbance was measured at a 450 nm wavelength in a Varioskan microtiter plate reader (ThermoFisher). Samples were run in duplicate and compared with the Glutamate standard after subtracting the blank lacking the Enzyme Mix.

### 2.14 Intracellular Ca^2+^ imaging

Two days before live cell imaging, confluent astrocytes were seeded on Matrigel^®^-coated 24-mm round glass coverslips in a 1:3 ratio. On the day of imaging, cells were treated with a loading solution of DMEM/F-12 w/o Phenol red (Gibco^™^) containing 2 μM cell-permeant Fluo-4 acetoxymethyl ester (Fluo-4-AM, Thermo Fisher) with the addition of the equal quantity (1:1 v/v) of 20% Pluronic F-127 (Thermo Fisher), to assist in the dispersion of the nonpolar AM ester in the aqueous medium. After incubation with the loading solution for 30 min at 37 °C, the astrocytes were washed thoroughly three times with DMEM/F-12 w/o Phenol red to remove any dye non-specifically associated with the cell surface. Finally, coverslips were mounted in live-cell imaging chambers and immediately transferred to a Leica SP5 AOBS inverted confocal microscope equipped with a live-cell module maintaining a 37 °C, 5% CO_2_, and >90% relative humidity environment. After a 1-min time-lapse series acquisition, ATP solution (Sigma-Aldrich) was added to the medium. The final concentration of ATP in the medium was 100 μM. The ATP was added carefully using a conventional pipette, without touching the imaging chamber, to avoid movement artifacts and ensure that the same field of view was imaged before and after the ATP application. Finally, a 10-min time-lapse series were acquired to record the ATP-induced fluorescence. Series were recorded at 400 Hz with 1 frame per second (fps) acquisition. Each frame was detected on a spectral PMT detector with an HCX PL APO CS 20×/0.7 DRY UV lens. Cells were irradiated with the argon laser.

### 2.15 Quantification of calcium imaging traces with deCLUTTER^2+^ pipeline

All images were processed using Fiji software (version 1.53c). Raw time-lapse imaging stacks (*x, y, t*) of intracellular Ca^2+^ were first corrected for a drift using a *Correct 3D Drift* plug-in (Parslow et al., 2014). To obtain individual Ca^2+^ traces, cells were segmented with a semi-automated segmentation strategy adapted from a method described by Schwendy et al (Schwendy et al., 2019). To estimate cell borders, the local maxima (mainly located in the cell nucleus) of each cell in a Z-projected image were determined to create an inverted tile mask with one segmented particle (tile) per maximum. Next, another mask was made using a Li background threshold method to select the total cell area in the image. In this case, the “leaky” fluorescence from the FLUO-4-AM was exploited to identify all the cell bodies via a thresholding method. A logical *XOR* operation using an *Image Calculator* was performed on both masks. To smooth the objects and remove isolated pixels, erosion and dilation were performed using an *Open* function in the Binary submenu. The remaining holes were filled using a *Fill Holes* function in the Binary submenu. Regions of Interest (ROIs) were created using an *Analyze Particle* plug-in. Each ROI was manually inspected to ensure that it defines only one cell. ROIs connected at one or two pixels were separated with the *Selection Brush*.Lastly, mean grey values per time frame were calculated in each ROI. For each time point in each ROI, we calculated the signal-to-baseline ratio of fluorescence F using the formula 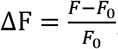, where the baseline F0 is estimated as the average of the fluorescence levels of 20 time points before ATP addition. The traces were visualized using a ggplot2 v3.3.6 and pheatmap v1.0.12 R packages.

To capture the time functions underlying the calcium influx behaviors observed in the astrocytes, we employed functional principal component analysis (fPCA). With this technique, it is possible to analyze a set of observations ordered in time, i.e., functions, and to identify the underlying eigenfunctions (ϕ) that describe the shape of the data. Similarly to the principal components in PCA, ϕ are ranked by the amount of variance they explain and can be used to reduce the dimensionality of the dataset. ΔF/F0 values from the six cell lines were merged and analyzed with the fdapace v0.5.9 (Jane-Ling et al., 2016) R package. Next, we selected the top ϕs explaining 95% of the variance in the data. We used them as an input for uniform manifold approximation and projection (UMAP) onto two dimensions using the umap v0.2.9.0 R package. Finally, we performed k-means clustering using the factoextra v1.0.7 0 R package and selected k = 3 using the elbow plot method and visual evaluation of the UMAP plot.

### 2.16 Statistical analyses

Statistical analyses were carried out using GraphPad Prism 9 (San Diego, USA). Unpaired t-test was performed for the analysis of the glutamate uptake assay. Two-way ANOVA with post hoc Tukey test were applied to the immunocytochemistry of astrocytes. Effects with p-value < 0.05 were considered significant.

## 3. Results

### 3.1 Astrocytes are derived from ventral midbrain patterned progenitor cells

We developed a strategy to differentiate human-derived iPSCs into astrocytes by combining various steps from previously described astrocyte differentiation protocols and the principles of astrocyte development in vivo. First, we generated eight iPSC lines that formed compact colonies with well-defined edges and expressed pluripotency markers, including NANOG, SSEA-4, OCT-4, and TRA-1-81 (**Supplementary Fig. 1A**). Next, we generated neural progenitor cells patterned towards a ventral midbrain identity that are able to also differentiate into dopaminergic neurons (**Supplementary Fig. 1B**, **1C**, and **1D**) as previously described (Grochowska et al., 2021; Monzel et al., 2017; Quadri et al., 2018; Reinhardt et al., 2013; Vanhauwaert et al., 2017). These progenitor lines expressed neural progenitor markers such as SOX2 and nestin but remained undifferentiated, as demonstrated by low expression of the mature neuron marker MAP2 (**Supplementary Fig. 1D**).

In vivo, neural progenitors undergo many rounds of neurogenesis before committing to the glial fate (Miller and Gauthier, 2007). Hence, we strived to accelerate gliogenesis in vitro by combining several well-established approaches. It has been shown that culturing astrocytes in a three-dimensional matrix induces the expression of astrocyte-specific genes and improves astrocyte maturity (Lattke et al., 2021). Therefore, we aggregated progenitor cells to form floating spheres (**Fig. 1A, 1Bb, and 1Bc**). Next, we exposed the spheres to basic fibroblast growth factor (bFGF) and epidermal growth factor (EGF) to mass amplify progenitor cells. Consecutively, we added leukemia inhibitory factor (LIF) and EGF to accelerate glial differentiation by the activation of the JAK-STAT signaling pathway (Bonni et al., 1997; Perriot et al., 2018) (**Fig. 1A**). Finally, when astrocyte-enriched spheres were plated, astrocytes migrated radially out of the spheres in the presence of ciliary neurotrophic factor (CNTF) for further maturation (**Fig. 1A** and **1Bd**). Out of eight cell lines, two lines failed to attach firmly to the plates. Therefore, they were excluded from further analyses. This procedure led to a proliferative population of astrocytes expressing key astrocyte markers, including glial fibrillary acidic protein (GFAP), aquaporin 4 (AQP4), astrocytic precursor markers CD44, and nestin in all human iPSC-derived astrocytes as demonstrated by ICC (**Fig. 1C**). On average, this protocol yielded 82% of GFAP-expressing astrocytes at week 13 (**Fig. 1D**). The GFAP expression increased to 90% after 20 weeks in culture (**Fig. 1E**). At both time points, there were significant differences detected in cell composition between lines (**Fig. 1D** and **1E**). No evident contamination by neurons was found, as assessed by immunostaining with an anti-MAP2 antibody, confirming the efficiency of this protocol in generating highly enriched astrocyte populations (**Fig. 1C-E**).

**Fig. 1.**
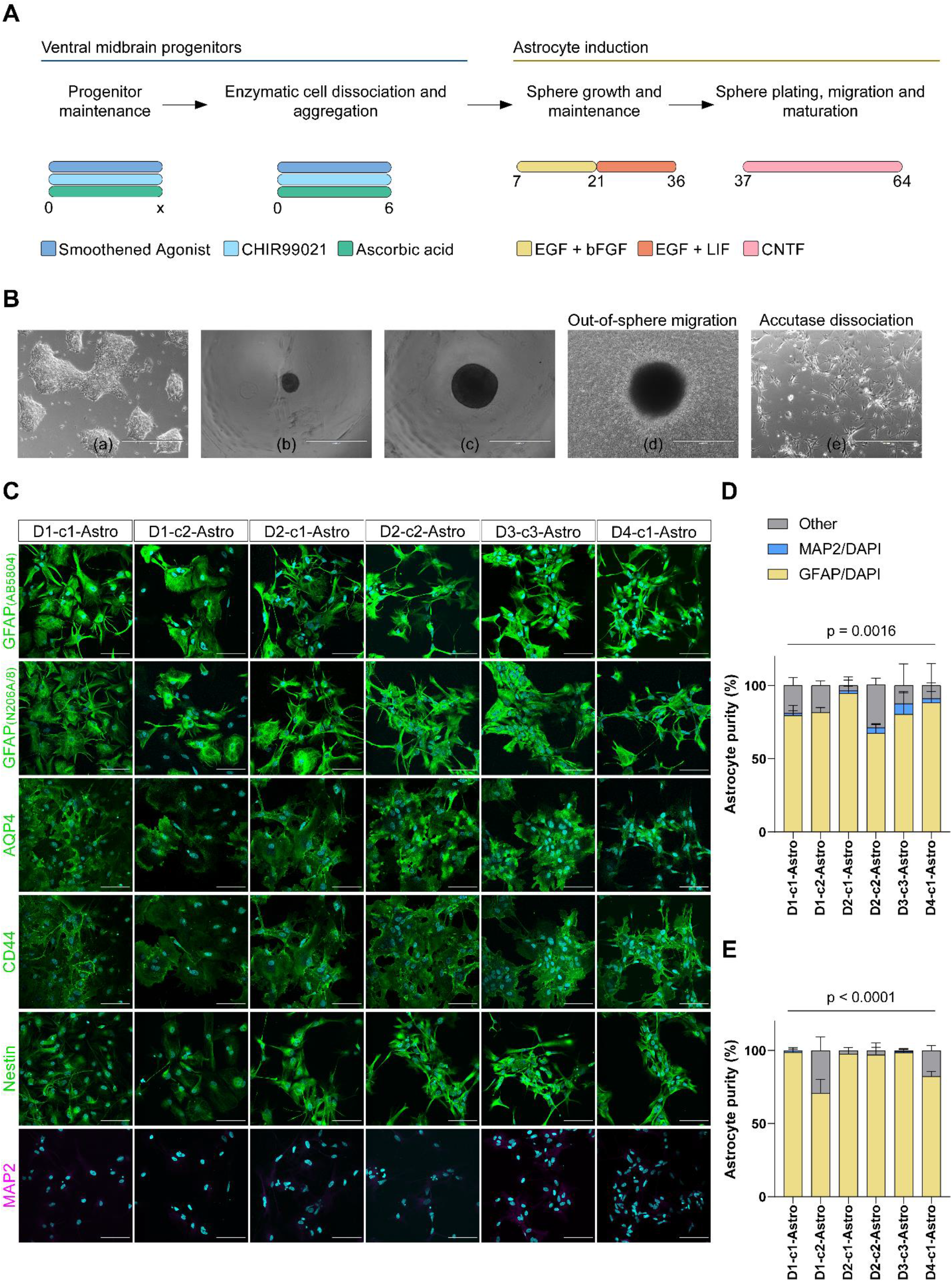
Generation and immunocytochemical characterization of astrocytes derived from ventral midbrain patterned progenitors. **A.** Schematic of the astrocyte differentiation protocol depicting the major steps with the accompanying supplements. **B.** Representative bright field images of various cell morphologies at different culturing steps during differentiation. Ventral midbrain progenitors (a) were seeded in low attachment plates to form spheres (b). Spheres were cultured for several weeks in EGF/bFGF or EGF/LIF-containing media (c). Spheres were plated and further differentiated with CNTF (d). Once matured, astrocytes were enzymatically dissociated and used for downstream assays (e). **C.** Representative ICC images of 20 weeks old astrocytes from six lines staining positive for general astrocyte markers GFAP and AQP4, for astrocytic precursor markers CD44 and nestin, and staining negative for the mature neuron marker MAP2. Scale bars, 100 μm. Nuclei were counterstained with DAPI (cyan). Astrocyte cultures mainly comprise GFAP-positive cells at 13 (**D**) and 20 weeks (**E**). The number of independent differentiations per line = 2. Two-way ANOVA, interaction *p*-value

### 3.2 Distinct expression profiles separate precursors and astrocytes

To investigate the molecular characteristics and whether the observed morphological changes were consistent with changes in the transcriptional landscape, we sequenced RNA from eight progenitor and six astrocyte lines. First, we employed eSNP-Karyotyping on the RNA-seq data to check for chromosomal aberrations. No large chromosomal multiplication could be detected in any of the progenitor lines (**Supplementary Fig. 2**). eSNP-Karyotyping revealed a chr2 multiplication in one of the astrocyte lines. Other astrocyte lines showed patterns similar to the corresponding precursor lines. Hence, we excluded the D4-c1 astrocyte line from the following transcriptomic analyses.

Next, we aimed to determine whether expression profiling could unbiasedly stratify the generated cell lines. PCA analysis on the top 500 most variable genes revealed two well-distinct groups corresponding to the two populations. PC1, capturing 82% of the variability in gene expression, clearly discriminated between the two (**Fig. 2A**).

**Fig. 2.**
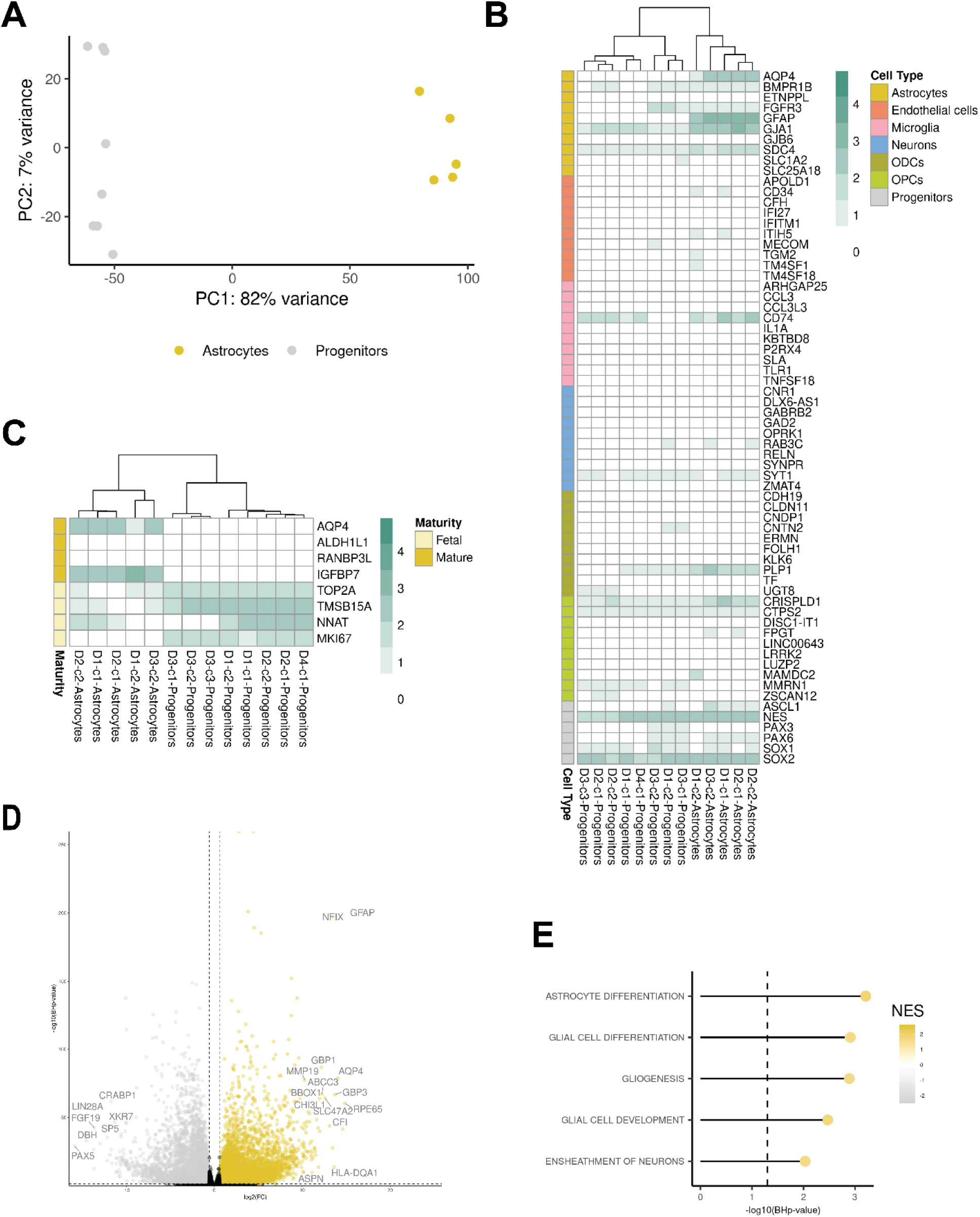
Transcriptomic characterization of progenitors and astrocytes. **A.** PCA plot showing line clustering conducted on top 500 variable genes. **B.** Heatmap showing the level of expression as log10(FPKM+1) of top 10 BRETIGEA and literature-described markers for brain cell types. **C.** Heatmap showing the level of expression as log_10_(FPKM+1) of literature-derived fetal and mature astrocyte markers. **D.** Volcano plot showing significantly upregulated (yellow) and downregulated (grey) genes in the astrocyte lines compared to the progenitor lines. **E.** Lollipop plot showing top 5 (ordered by *BHp-*value) significantly enriched gene ontology biological processes among the nervous system development (GO:0007399) child terms

We assessed the expression of the top 10 markers selected from BRETIGEA encompassing six cell types (astrocytes, endothelial cells, neurons, microglia, oligodendrocytes, and oligodendrocytes precursor cells) and from literature for the progenitors to identify the cell lines. The precursors and the astrocytes expressed the expected cell-type specific markers consistently. In contrast, lower-to-no expression of the majority of markers of the other cell types could be detected (**Fig. 2B**). However, the astrocytes expressed some of the markers of the precursor cells. For this reason, we assessed the maturity level of the generated astrocytes using a list of markers for fetal and mature astrocytes (Sloan et al., 2017) (**Fig. 2C**). Compared to the precursor cells, astrocytes expressed a high level of two of the mature astrocyte markers (*AQP4* and *IGFBP7*). Lower expression levels of fetal astrocytes markers could still be detected in the astrocytes.

### 3.4 Genome-wide expression profiling confirms effective differentiation

We performed differential expression analysis on the RNA-seq data to identify the genes driving the differences between the precursors and astrocytes we generated. A total of 13,452 DEGs was detected. 7,196 were upregulated in the astrocytes compared to the progenitors (**Fig. 2D**, **Supplementary Table 6**). Among the top up-regulated genes, we found established astrocytic markers, including *GFAP* (log2FC 16.87 ± 0.55, *BHp*-value 3.84 × 10^-201^) and *AQP4* (log2FC 14.07 ± 0.73, *BHp*-value 4.02 × 10^-79^). Similarly, among the top down-regulated genes, we identified genes related to the stemness and proliferative potential of the precursor cells, including *LIN28A* (log2FC −16.51 ± 1.02, *BHp*-value 3.59 × 10^-57^) and *PAX5* (log2FC −15.88 ± 1.37, *BHp*-value 1.07 × 10^-29^).

We also assessed the effect of the differentiation at the pathway level by performing GSEA on the genes ranked by the differential expression analysis statistics. 1,282 terms from the gene ontology biological process category were significantly different between the two cell types, with 808 being upregulated in the astrocytes confirming broad differences in the transcriptomes of the two lines (**Supplementary Table 7**). We prioritized the nervous system development (GO:0007399) child terms from this category to focus on the differentiation effect. Of the 181 terms passing our filtering criteria, 16 significantly differed between astrocytes and precursors. The top significantly up-regulated term was astrocyte differentiation (NES = 1.86, *BHp-value* 6.16 × 10^-4^), further supporting the directionality and effectiveness of the differentiation protocol (**Fig. 2E, Supplementary Table 7**).

### 3.5 Astrocytes respond to pro-inflammatory cytokines and display a low extracellular glutamate uptake

Reactive astrocytes play an important role in the pathogenesis of many neurodegenerative diseases (Escartin et al., 2021). To investigate this, several studies have modeled inflammation-stimulated reactivity in iPSC-derived astrocytes (Barbar et al., 2020; Leng et al., 2022; Perriot et al., 2018; Roybon et al., 2013; Santos et al., 2017; Tchieu et al., 2019; Tcw et al., 2017). We also characterized the transcriptomic profile of our astrocytes stimulated with pro-inflammatory cytokines TNF-α, IL-1α, and C1q that drive an A1-like reactive state (Liddelow et al., 2017) using qPCR. iPSC-derived astrocytes exhibited morphological changes upon the cytokine treatment, including the remodeling of GFAP-positive intermediate filaments (**Fig. 3A**). A proportion of treated astrocytes appeared to have more cellular processes containing GFAP intermediate filaments and more intermediate filament branches than the untreated astrocytes (**Fig. 3A**, boxed areas). Next, we assessed the expression of *C3*, *LCN2, SERPINA3*, and *GFAP* transcripts affected in astrocytes in vivo and in vitro upon A1 stimulation (Barbar et al., 2020; Liddelow et al., 2017). TNF-α, IL-1α, and C1q treated astrocytes showed strong upregulation of *C3*, *LCN2*, and *SERPINA3* mRNA transcripts as assessed by the RT-qPCR (**Fig. 3B-D**). The levels of upregulation of *C3, LCN2*, and *SERPINA3* showed heterogeneous responses across the different astrocyte lines. Strikingly, *GFAP* expression was downregulated in the treated astrocytes across all cell lines (**Fig. 3E**). This contrasts with the changes in the astrocytic cytoskeleton and hypertrophy we detected with GFAP immunocytochemistry in the treated astrocytes. Overall, we observed similar expression patterns to previous in vitro and in vivo studies (Barbar et al., 2020; Liddelow et al., 2017). These results show that the generated astrocytes are immunocompetent and respond to inflammatory stimuli by changing their morphology and transcript expression.

**Fig. 3.**
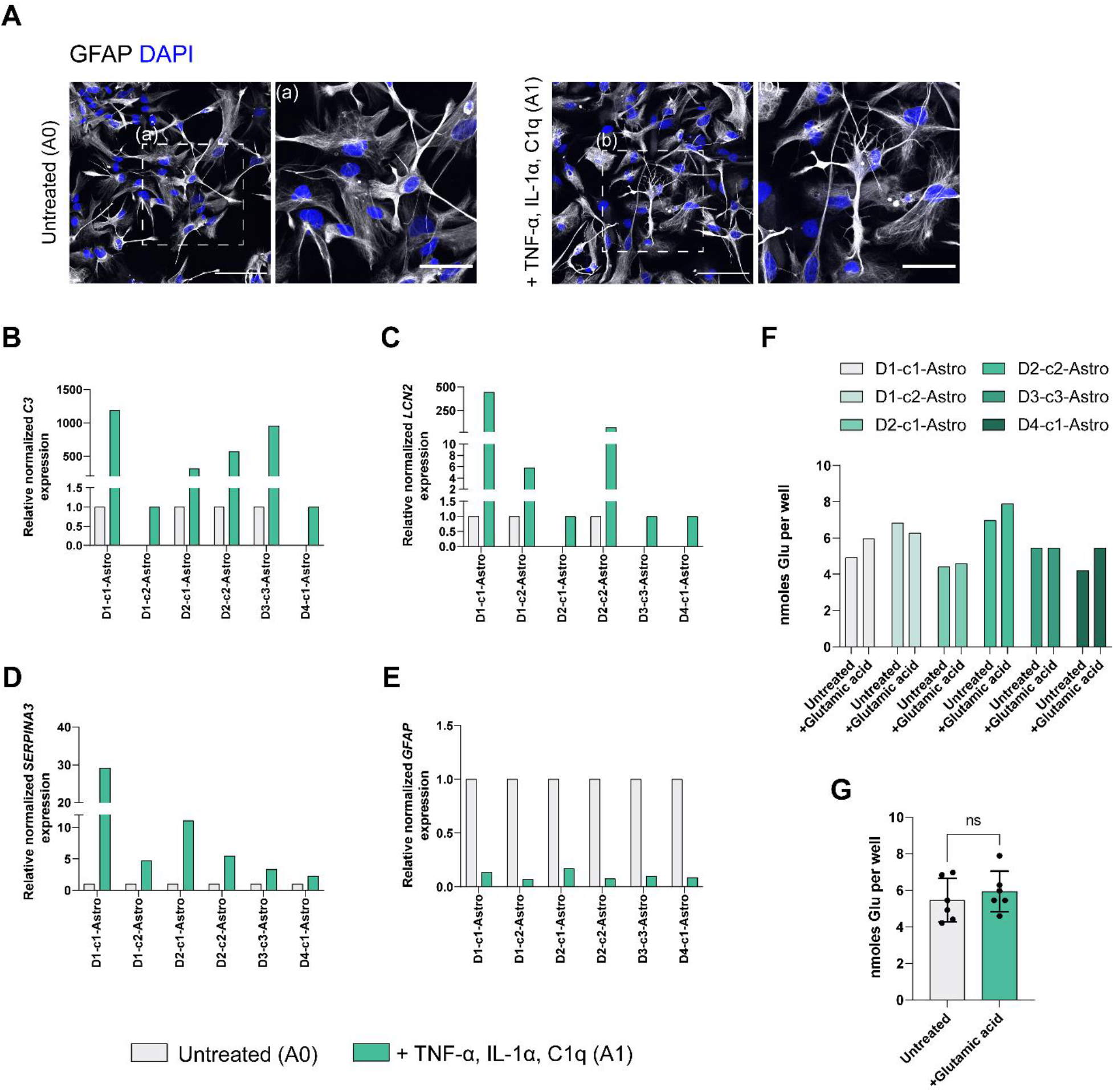
Functional characterization of astrocytes. **A.** Representative ICC images showing GFAP (grey) expression in astrocytes upon stimulation with TNF-α, IL-1α, and C1q. Nuclei were counterstained with DAPI (blue). White dashed boxes indicate the magnified images (a, b) to highlight changes in morphology. Scale bars, 100 μm and 50 μm. **B-E**. 20 weeks old astrocytes upregulate reactive astrocyte-markers *C3* (**B**), *LCN2* (**C**), *SERPINA3* (**D**), and *GFAP* (**E**) upon TNF-α, IL-1α, and C1q treatment. Bar charts depict RT-qPCR analysis. Data represent the mean of relative normalized mRNA expression from three technical replicates. *CLK2, COPS5*, and *RNF10* were used as housekeeping genes. **F**. Glutamate uptake analysis on 20 weeks astrocytes. Bar graphs show nmoles of glutamate taken up by each astrocyte line after incubation with 100 μM glutamate for 3 h compared with untreated wells. Data represent the mean nmoles from two technical replicates. **G.** Pooled results from **F**. Each dot represents an astrocyte line. Unpaired t-test; ns, not significant

We next tested another important astrocyte physiological function, the uptake of glutamate, which prevents neuronal excitotoxicity in vivo (Verkhratsky and Nedergaard, 2018). All lines presented detectable levels of intracellular glutamate in baseline conditions (average 5.47 ± 1.195 nmol). Five out of six astrocyte lines showed a slight increase in the uptake of extracellular glutamate upon 3 h treatment (average 5.94 ± 1.113 nmol) (**Fig. 3F**). However, these findings were not significant in pooled testing (**3G**).

### 3.6 Fluoro-4-AM imaging confirms electrophysiological responsiveness of astrocytes and deCLUTTER^2+^pipeline reveals dominant profiles and clusters in calcium traces

Neurotransmitter-induced intracellular Ca^2+^ transients play a pivotal role in astrocyte functionality and have been observed in vivo and ex vivo systems (Gorzo and Gordon, 2022; Lia et al., 2021). We thus wanted to assess whether the generated astrocytes possessed a similar physiological property.

We used Fluo-4-AM cell loading coupled with ATP stimulation to analyze the calcium dynamics in the astrocyte lines. We imaged all the cell lines and identified green-fluorescent ROIs in the somata using Fiji. We selected 52 cells per line, resulting in a total of 312 cells across six cell lines. Heatmaps showed that the different cell lines had a heterogeneous response to the ATP stimulus with a variable number of responsive cells. These cells were characterized by distinct patterns with an average peak amplitude of 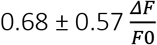 (**Fig. 4A**). Although to different extents, an increase in fluorescence upon the ATP stimulus was evident in all cell lines.

**Fig. 4.**
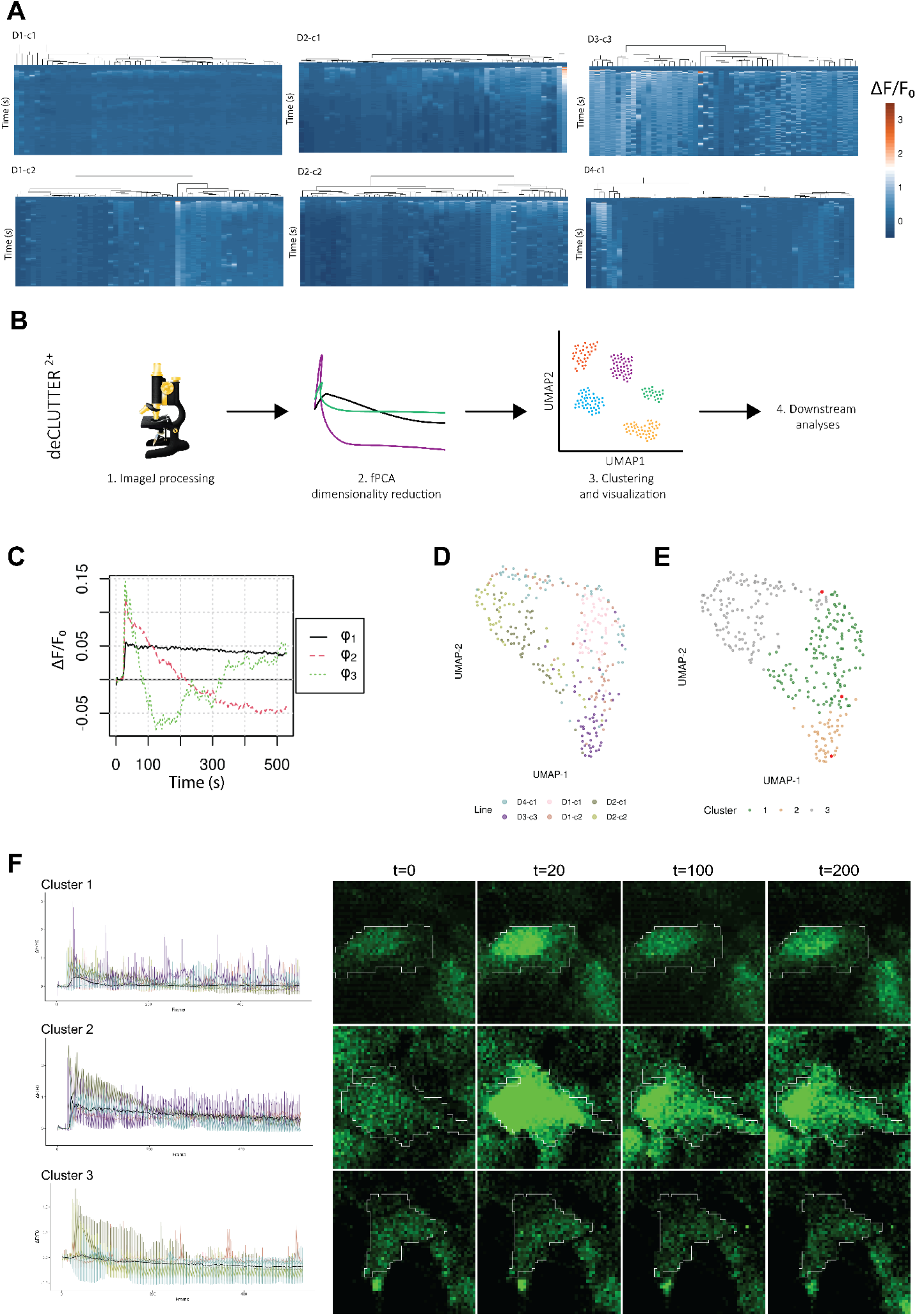
Characterization of ATP-induced Ca^2+^ transients in astrocytes. **A.** Heatmaps showing normalized ΔF/F_0_ values for the randomly selected cells for each cell line (n = 52). **B**. Schematic overview of deCLUTTER^2+^pipeline. **C.** fPCA top 3 eigenfunctions extracted from the ΔF/F0 across the cell lines. **D.** UMAP plot showing astrocyte lines spread. **E.** UMAP plot showing clustering of the cells into the three k-means clusters. **F.** ΔF/F0 profiles of the three defined clusters along the imaging time course. Tracks are colored by cell line, and the median ΔF/F0 is highlighted in black. Representative images of the three clusters sampled at time points t=0, 20, 100, 200 s. The randomly selected cells are highlighted in red in Fig. 4D

To further characterize the ATP-response behavior, we developed a novel pipeline we named deCLUTTER^2+^that takes as input the Fluo-4-AM signal calculated with Fiji across the cells. Briefly, fPCA is applied to perform dimension reduction and denoising (or deCLUTTER^2+^) the signals. Next, clusters are identified using k-means. Finally, UMAP is applied for visualization of the clusters (**Fig. 4B**).

Specifically, we applied fPCA to extract the main ϕs describing the dominant patterns of variability across cell lines. The top 3 ϕs explained 83% of the variability in the Ca^2+^ traces (**Fig. 4C**). ϕ_1_ (68% of the variability) captured part of the stimulus-associated increase in fluorescence and then it remained nearly constant with just a slow decrease. ϕ_2_ (12% of the variability) encoded the sharp variation in fluorescence around the ATP stimulus time point in responsive cells. Finally, ϕ_3_ (2.6% of the variability) captured both the variability around the ATP stimulus time point and another increase in fluorescence at about half the imaging time course. To show the local and global structures in the cell lines with a 2D representation, we constructed a lower dimensionality embedding with UMAP. We selected the top 22 ϕs, explaining 95% of the variance in the Fluo-4-AM tracks. The different cell lines were admixed onto the UMAP plot, showing a continuum with regions of higher density (**Fig. 4D**). In the case of the D2-derived lines, it was possible to see an overlap that might be partially driven by a donor effect. We clustered the cell lines using k-means clustering and obtained three distinct groups with a differential contribution from the various cell lines (**Fig.4E**). The three clusters were characterized by different 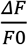 profiles. The cell lines contributed a variable number of cells to each cluster (**Fig. 4F**). Clusters 1 and 3 corresponded to low and non-responsive cells, whereas cluster 2 contained cells highly responsive to the ATP stimulus.

## 4. Discussion

iPSC-derived modeling is gaining momentum as a complementary method to study the properties of human-derived cells under physiological and pathological conditions. However, specific protocols for the generation of iPSC-derived ventral midbrain astrocytes to study the potential role of astrocytes in the non-cell autonomous neurodegeneration observed in PD, have been lagging, and their morphological and functional characterization are limited (**Supplementary Table 1**). In our study, we addressed some of these limitations and generate midbrain-specified astrocytes by exploiting developmental events required for astrogenesis.

Recent data indicates that ventral midbrain astrocytes differ from telencephalic astrocytes at the physiological and transcriptomic levels (Siletti et al., 2022; Xin et al., 2019). However, what could be driving these differences remains to be investigated. Several studies have proposed that such regional heterogeneity of astrocytes arises early in development. The patterning along the neuraxis by sonic hedgehog (SHH), fibroblast growth factors (FGFs), WNTs, and bone morphogenic factors (BMPs) leads to a rostral-caudal and dorsal-ventral segmentation of the neuroepithelium into domains that give rise to distinct subtypes of progenitors encoding regional information (Hochstim et al., 2008; Rowitch and Kriegstein, 2010; Sardar et al., 2020; Tsai et al., 2012). Studies performed in various brain regions have demonstrated that astrocytes and neurons derived from a common progenitor show a shared region-specific molecular signature, which might act as a code for region-specific interactions (Herrero-Navarro et al., 2021). Moreover, astrocytes and neurons derived from a common progenitor domain migrate radially and share their final position (Herrero-Navarro et al., 2021; Torigoe et al., 2015). Hence, leveraging the same progenitor pool for the generation of astrocytes and neurons in vitro might enable studying neuron-astrocyte interactions of cells carrying a region-specific profile.

Although the developmental trajectory of the ventral midbrain astrocytes is poorly defined, we speculated that the early patterning step would partially recapitulate the ventral midbrain ontogeny. Therefore, we leveraged well-described patterning protocols using morphogens that specify the ventral midbrain identity (Reinhardt et al., 2013). Importantly, the generated progenitor pool can also differentiate into vmDANs (Grochowska et al., 2021; Reinhardt et al., 2013; Vanhauwaert et al., 2017). Our protocol resulted in a highly pure population of cells expressing classical astrocyte markers. Furthermore, our astrocytes showed strong transcriptomic evidence of the expected differentiation and expressed mature-astrocyte markers, including AQP4, GFAP, and Connexin 43/GJA1. However, the low expression of some fetal-astrocyte markers and the inefficient extracellular glutamate uptake suggest that subpopulations of lesser differentiated astrocytes co-exist with more mature ones. On the other hand, the results of the glutamate uptake assay might have been influenced by the presence of glutamate in the baseline culturing media warranting caution in the interpretation of the data. The ratio of mature to immature cells might be further increased using maturation-accelerating supplements like the ones reported in a recent preprint for iPSC-derived neurons (Hergenreder et al., 2022). Unfortunately, the absence of a specific midbrain-astrocyte marker makes it challenging to validate the ontogeny of the generated cell line. However, our astrocytes expressed high levels of GFAP, in keeping with its high expression in subcortical regions, while lower expression is reported in the cortical regions and the cerebellum (Siletti et al., 2022; Torres-Platas et al., 2016).

Genomic instability can have profound effects on cellular phenotypes, including growth speed, differentiation potential, and resistance to stress (Weissbein et al., 2016). Therefore, we employed eSNP-Karyotyping on the RNA-seq of all the cell lines and identified chromosomal aberrations in one of the astrocyte lines (**Supplementary Fig. 2**). This highlights the importance of assessing for culturing-induced chromosomal aberrations not only when iPSCs are generated but also after differentiation. This aspect has surprisingly received little attention in previous literature despite the consequences that such events may have on downstream analyses.

Next, we aimed to generate a model that recapitulates some of the functional properties of astrocytes. We emphasize our findings from these functional assays, establishing iPSC-derived astrocyte cultures as a platform for disease modeling. Recent research using post-mortem samples identified reactive astrocytes in the midbrain of individuals with PD (Schonhoff et al., 2020). Therefore, we investigated the responses of astrocytes to pro-inflammatory stimuli by treating the cells with TNF-α, IL-1α, and C1q that drive a reactive phenotype (Escartin et al., 2021; Liddelow et al., 2017). This assay has been widely used to study neuroinflammation in vitro and in vivo (Barbar et al., 2020; Leng et al., 2022; Liddelow et al., 2017; Zhou et al., 2019). Moreover, the morphology and transcriptomic profile of stimulated astrocytes have been well characterized. Our protocol generated astrocytes that were immunocompetent. In particular, we observed a strong upregulation of complement component 3 (C3), also found in the astrocytes residing in the substantia nigra pars compacta (SNpc) of idiopathic PD cases (Liddelow et al., 2017). Moreover, we observed similar expression patterns for *GFAP, LCN2*, and *SERPINA3* transcripts, which were in line with in vivo studies and previously published iPSC-derived astrocyte protocols (Barbar et al., 2020; Liddelow et al., 2017). Altogether, we characterized a cellular in vitro model that has the potential to be applied in mechanistic studies on neuroinflammatory signaling in PD.

The effectiveness of our protocol is further supported by the data showing heterogeneous Ca^2+^ transients in ventral midbrain astrocytes upon ATP treatment. These currents are known to have a critical impact on neurotransmitter release (Bezzi and Volterra, 2001), synaptic transmission and plasticity (Di Castro et al., 2011; Panatier et al., 2011), and even behavior and cognition (Kofuji and Araque, 2021; Nagai et al., 2021; Santello et al., 2019) and are widely used as functional readouts in vivo and in vitro (Gorzo and Gordon, 2022; Lia et al., 2021). Importantly, calcium dysregulation is a potential mechanism for several genes implicated in PD (Zaichick et al., 2017). Abnormal calcium signaling in the astrocytes might cause dysfunction in dopaminergic neurons, activate microglia, and disrupt the blood brain barrier integrity, therefore contributing to the pathological mechanisms seen in PD (Bancroft and Srinivasan, 2021). The classical processing of calcium images from iPSC-derived astrocytes has encompassed visualization of the transients via time-lapse imaging and traces. To better integrate the data from the different cell lines, we developed the novel deCLUTTER^2+^ pipeline. deCLUTTER^2+^ can be used to perform semiautomatic recognition of spontaneous or cue-dependent recurring patterns in fluorescence time-lapse imaging. Our approach can handle variability and can be used to easily visualize local and global structures among the analyzed cells. We anticipate that this approach could be used in the context of disease modeling to study the association of specific cluster(s) to conditions.

In conclusion, we have generated ventral midbrain astrocytes that express astrocyte-specific markers and recapitulate some of the physiological properties of in vivo ventral midbrain astrocytes. Our ventral midbrain astrocyte model can also be integrated with neurons and other glial cell types, going beyond traditional 2D co-culture systems (Majumdar et al., 2011; Park et al., 2018). Such neuron-glia interaction paradigms might help elucidate the pathological processes observed in neurodegenerative diseases, for example PD, and potentially pave the way toward disease-modifying treatments.

## Supporting information

Supplementary Table6_DEA

Supplementary Table7_GSEA

## Supplementary Material

**Supplementary Fig. 1** Generation and characterization of iPSCs and iPSC-derived ventral midbrain patterned progenitors

**Supplementary Fig. 2** eSNP-Karyotyping reveals chromosomal multiplication in one astrocyte line

**Supplementary Table 1** Differentiation protocols that generate ventral midbrain astrocytes

**Supplementary Table 2** Cell culture media composition

**Supplementary Table 3** Details of the primary antibodies used in this study

**Supplementary Table 4** Details of the 14 stem cell-derived clonal cell lines used in this study

**Supplementary Table 5** qPCR primers

**Supplementary Table 6** Differential expression analysis results report

**Supplementary Table 7** Gene set enrichment analysis results report

## Acknowledgments

We thank Erasmus MC iPS Core Facility for providing the hiPSC lines and the Erasmus MC Optical Imaging Center (OIC) members for their microscopy assistance.

## Author contributions

M.M.G., F.F., D.N., and W.M. designed the experiments. M.M.G., A.C.M., D.N., Va.B., and G.J.B. performed the experiments. M.M.G., F.F., A.C.M., and D.N. performed data analysis. M.M.G., F.F., A.C.M., D.N., V.B., and W.M interpreted the data. M.M.G. and F.F. wrote the initial draft of the manuscript. W.M. and V.B. conceived the study and supervised the experiments. All authors critically read the manuscript and approved the final version.

## Data Availability Statement

All analyses were conducted in R version v4.2.1 and ImageJ version 1.53c (Java 1.8.0_172 64-bit). Datasets are available upon reasonable request.

## References

Andrews, S. (2010). FASTQC. A quality control tool for high throughput sequence data.

Bancroft, E. A. and Srinivasan, R. (2021). Emerging Roles for Aberrant Astrocytic Calcium Signals in Parkinson’s Disease. Front Physiol 12, 812212.

Bandopadhyay, R., Kingsbury, A. E., Cookson, M. R., Reid, A. R., Evans, I. M., Hope, A. D., Pittman, A. M., Lashley, T., Canet-Aviles, R., Miller, D. W. et al. (2004a). The expression of DJ-1 (PARK7) in normal human CNS and idiopathic Parkinson’s disease. Brain 127, 420–30.

Bandopadhyay, R., Kingsbury, A. E., Cookson, M. R., Reid, A. R., Evans, I. M., Hope, A. D., Pittman, A. M., Lashley, T., Canet-Aviles, R., Miller, D. W. et al. (2004b). The expression of DJ-1 (PARK7) in normal human CNS and idiopathic Parkinson’s disease. Brain 127, 420–430.

Barbar, L., Jain, T., Zimmer, M., Kruglikov, I., Sadick, J. S., Wang, M., Kalpana, K., Rose, I. V. L., Burstein, S. R., Rusielewicz, T. et al. (2020). CD49f Is a Novel Marker of Functional and Reactive Human iPSC-Derived Astrocytes. Neuron 107, 436–453 e12.

Batiuk, M. Y., Martirosyan, A., Wahis, J., de Vin, F., Marneffe, C., Kusserow, C., Koeppen, J., Viana, J. F., Oliveira, J. F., Voet, T. et al. (2020). Identification of region-specific astrocyte subtypes at single cell resolution. Nat Commun 11, 1220.

Bayraktar, O. A., Bartels, T., Holmqvist, S., Kleshchevnikov, V., Martirosyan, A., Polioudakis, D., Ben Haim, L., Young, A. M. H., Batiuk, M. Y., Prakash, K. et al. (2020). Astrocyte layers in the mammalian cerebral cortex revealed by a single-cell in situ transcriptomic map. Nat Neurosci 23, 500–509.

Beard, E., Lengacher, S., Dias, S., Magistretti, P. J. and Finsterwald, C. (2021). Astrocytes as Key Regulators of Brain Energy Metabolism: New Therapeutic Perspectives. Front Physiol 12, 825816.

Bezzi, P. and Volterra, A. (2001). A neuron-glia signalling network in the active brain. Curr Opin Neurobiol 11, 387–94.

Bonni, A., Sun, Y., Nadal-Vicens, M., Bhatt, A., Frank, D. A., Rozovsky, I., Stahl, N., Yancopoulos, G. and Greenberg, M. E. D. (1997). Regulation of gliogenesis in the central nervous system by the JAK-STAT signaling pathway. Science 278, 477–83.

Booth, H. D. E., Hirst, W. D. and Wade-Martins, R. (2017). The Role of Astrocyte Dysfunction in Parkinson’s Disease Pathogenesis. Trends Neurosci 40, 358–370.

Bugiani, M., Plug, B. C., Man, J. H. K., Breur, M. and van der Knaap, M. S. (2022). Heterogeneity of white matter astrocytes in the human brain. Acta Neuropathol 143, 159–177.

Chung, W. S., Allen, N. J. and Eroglu, C. (2015). Astrocytes Control Synapse Formation, Function, and Elimination. Cold Spring Harb Perspect Biol 7, a020370.

Dezonne, R. S., Stipursky, J., Araujo, A. P., Nones, J., Pavao, M. S., Porcionatto, M. and Gomes, F. C. (2013). Thyroid hormone treated astrocytes induce maturation of cerebral cortical neurons through modulation of proteoglycan levels. Front Cell Neurosci 7, 125.

Di Castro, M. A., Chuquet, J., Liaudet, N., Bhaukaurally, K., Santello, M., Bouvier, D., Tiret, P. and Volterra, A. (2011). Local Ca2+ detection and modulation of synaptic release by astrocytes. Nat Neurosci 14, 1276–84.

di Domenico, A., Carola, G., Calatayud, C., Pons-Espinal, M., Munoz, J. P., Richaud-Patin, Y., Fernandez-Carasa, I., Gut, M., Faella, A., Parameswaran, J. et al. (2019). Patient-Specific iPSC-Derived Astrocytes Contribute to Non-Cell-Autonomous Neurodegeneration in Parkinson’s Disease. Stem Cell Reports 12, 213–229.

Dobin, A., Davis, C. A., Schlesinger, F., Drenkow, J., Zaleski, C., Jha, S., Batut, P., Chaisson, M. and Gingeras, T. R. (2013). STAR: ultrafast universal RNA-seq aligner. Bioinformatics 29, 15–21.

Escartin, C., Galea, E., Lakatos, A., O’Callaghan, J. P., Petzold, G. C., Serrano-Pozo, A., Steinhäuser, C., Volterra, A., Carmignoto, G., Agarwal, A. et al. (2021). Reactive astrocyte nomenclature, definitions, and future directions. Nature Neuroscience 24, 312–325.

Ewels, P., Magnusson, M., Lundin, S. and Kaller, M. (2016). MultiQC: summarize analysis results for multiple tools and samples in a single report. Bioinformatics 32, 3047–8.

Gomes, F. C., Maia, C. G., de Menezes, J. R. and Neto, V. M. (1999). Cerebellar astrocytes treated by thyroid hormone modulate neuronal proliferation. Glia 25, 247–55.

Gomes, F. C., Sousa Vde, O. and Romao, L. (2005). Emerging roles for TGF-beta1 in nervous system development. Int J Dev Neurosci 23, 413–24.

Gonzalez-Reyes, R. E., Nava-Mesa, M. O., Vargas-Sanchez, K., Ariza-Salamanca, D. and Mora-Munoz, L. (2017). Involvement of Astrocytes in Alzheimer’s Disease from a Neuroinflammatory and Oxidative Stress Perspective. Front Mol Neurosci 10, 427.

Gorzo, K. A. and Gordon, G. R. (2022). Photonics tools begin to clarify astrocyte calcium transients. Neurophotonics 9, 021907.

Grochowska, M. M., Carreras Mascaro, A., Boumeester, V., Natale, D., Breedveld, G. J., Geut,H., van Cappellen, W. A., Boon, A. J. W., Kievit, A. J. A., Sammler, E. et al. (2021). LRP10 interacts with SORL1 in the intracellular vesicle trafficking pathway in non-neuronal brain cells and localises to Lewy bodies in Parkinson’s disease and dementia with Lewy bodies. Acta Neuropathol 142, 117–137.

Hergenreder, E., Zorina, Y., Zhao, Z., Munguba, H., Calder, E. L., Baggiolini, A., Minotti, A. P., Walsh, R. M., Liston, C., Levitz, J. et al. (2022). Combined small molecule treatment accelerates timing of maturation in human pluripotent stem cell-derived neurons. bioRxiv, 2022.06.02.494616.

Herrero-Navarro, A., Puche-Aroca, L., Moreno-Juan, V., Sempere-Ferrandez, A., Espinosa, A., Susin, R., Torres-Masjoan, L., Leyva-Diaz, E., Karow, M., Figueres-Onate, M. et al. (2021). Astrocytes and neurons share region-specific transcriptional signatures that confer regional identity to neuronal reprogramming. Sci Adv 7.

Hochstim, C., Deneen, B., Lukaszewicz, A., Zhou, Q. and Anderson, D. J. (2008). Identification of positionally distinct astrocyte subtypes whose identities are specified by a homeodomain code. Cell 133, 510–22.

Ioannou, M. S., Jackson, J., Sheu, S. H., Chang, C. L., Weigel, A. V., Liu, H., Pasolli, H. A., Xu, C. S., Pang, S., Matthies, D. et al. (2019). Neuron-Astrocyte Metabolic Coupling Protects against Activity-Induced Fatty Acid Toxicity. Cell 177, 1522–1535 e14.

Jane-Ling, W., Jeng-Min, C. and Hans-Georg, M. (2016). Functional Data Analysis. Annual Review of Statistics and Its Application 3, 257–295.

Jessen, N. A., Munk, A. S., Lundgaard, I. and Nedergaard, M. (2015). The Glymphatic System: A Beginner’s Guide. Neurochem Res 40, 2583–99.

Kim, J. M., Cha, S. H., Choi, Y. R., Jou, I., Joe, E. H. and Park, S. M. (2016). DJ-1 deficiency impairs glutamate uptake into astrocytes via the regulation of flotillin-1 and caveolin-1 expression. Sci Rep 6, 28823.

Kim, K. S., Kim, J. S., Park, J. Y., Suh, Y. H., Jou, I., Joe, E. H. and Park, S. M. (2013). DJ-1 associates with lipid rafts by palmitoylation and regulates lipid rafts-dependent endocytosis in astrocytes. Hum Mol Genet 22, 4805–17.

Kofuji, P. and Araque, A. (2021). Astrocytes and Behavior. Annu Rev Neurosci 44, 49–67.

Krencik, R., Weick, J. P., Liu, Y., Zhang, Z. J. and Zhang, S. C. (2011). Specification of transplantable astroglial subtypes from human pluripotent stem cells. Nat Biotechnol 29, 528–34.

Lanjewar, S. N. and Sloan, S. A. (2021). Growing Glia: Cultivating Human Stem Cell Models of Gliogenesis in Health and Disease. Front Cell Dev Biol 9, 649538.

Lattke, M., Goldstone, R., Ellis, J. K., Boeing, S., Jurado-Arjona, J., Marichal, N., MacRae, J. I., Berninger, B. and Guillemot, F. (2021). Extensive transcriptional and chromatin changes underlie astrocyte maturation in vivo and in culture. Nat Commun 12, 4335.

Leberbauer, C., Boulme, F., Unfried, G., Huber, J., Beug, H. and Mullner, E. W. (2005). Different steroids co-regulate long-term expansion versus terminal differentiation in primary human erythroid progenitors. Blood 105, 85–94.

Leng, K., Rose, I. V. L., Kim, H., Xia, W., Romero-Fernandez, W., Rooney, B., Koontz, M., Li, E., Ao, Y., Wang, S. et al. (2022). CRISPRi screens in human iPSC-derived astrocytes elucidate regulators of distinct inflammatory reactive states. Nat Neurosci.

Lia, A., Henriques, V. J., Zonta, M., Chiavegato, A., Carmignoto, G., Gomez-Gonzalo, M. and Losi, G. (2021). Calcium Signals in Astrocyte Microdomains, a Decade of Great Advances. Front Cell Neurosci 15, 673433.

Liao, Y., Smyth, G. K. and Shi, W. (2014). featureCounts: an efficient general purpose program for assigning sequence reads to genomic features. Bioinformatics 30, 923–30.

Liddelow, S. A., Guttenplan, K. A., Clarke, L. E., Bennett, F. C., Bohlen, C. J., Schirmer, L., Bennett, M. L., Munch, A. E., Chung, W. S., Peterson, T. C. et al. (2017). Neurotoxic reactive astrocytes are induced by activated microglia. Nature 541, 481–487.

Liu, L., Zhang, K., Sandoval, H., Yamamoto, S., Jaiswal, M., Sanz, E., Li, Z., Hui, J., Graham, B. H., Quintana, A. et al. (2015). Glial lipid droplets and ROS induced by mitochondrial defects promote neurodegeneration. Cell 160, 177–90.

Livak, K. J. and Schmittgen, T. D. (2001). Analysis of relative gene expression data using real-time quantitative PCR and the 2(-Delta Delta C(T)) Method. Methods 25, 402–8.

Love, M. I., Huber, W. and Anders, S. (2014). Moderated estimation of fold change and dispersion for RNA-seq data with DESeq2. Genome Biol 15, 550.

MacMahon Copas, A. N., McComish, S. F., Fletcher, J. M. and Caldwell, M. A. (2021). The Pathogenesis of Parkinson’s Disease: A Complex Interplay Between Astrocytes, Microglia, and T Lymphocytes? Front Neurol 12, 666737.

Majumdar, D., Gao, Y., Li, D. and Webb, D. J. (2011). Co-culture of neurons and glia in a novel microfluidic platform. J Neurosci Methods 196, 38–44.

McKenna, A., Hanna, M., Banks, E., Sivachenko, A., Cibulskis, K., Kernytsky, A., Garimella, K., Altshuler, D., Gabriel, S., Daly, M. et al. (2010). The Genome Analysis Toolkit: a MapReduce framework for analyzing next-generation DNA sequencing data. Genome Res 20, 1297–303.

McKenzie, A. T., Wang, M., Hauberg, M. E., Fullard, J. F., Kozlenkov, A., Keenan, A., Hurd, Y. L., Dracheva, S., Casaccia, P., Roussos, P. et al. (2018). Brain Cell Type Specific Gene Expression and Co-expression Network Architectures. Sci Rep 8, 8868.

Miller, F. D. and Gauthier, A. S. (2007). Timing is everything: making neurons versus glia in the developing cortex. Neuron 54, 357–69.

Miyazaki, I. and Asanuma, M. (2020). Neuron-Astrocyte Interactions in Parkinson’s Disease. Cells 9.

Monzel, A. S., Smits, L. M., Hemmer, K., Hachi, S., Moreno, E. L., van Wuellen, T., Jarazo, J., Walter, J., Bruggemann, I., Boussaad, I. et al. (2017). Derivation of Human Midbrain-Specific Organoids from Neuroepithelial Stem Cells. Stem Cell Reports 8, 1144–1154.

Nagai, J., Yu, X., Papouin, T., Cheong, E., Freeman, M. R., Monk, K. R., Hastings, M. H., Haydon, P. G., Rowitch, D., Shaham, S. et al. (2021). Behaviorally consequential astrocytic regulation of neural circuits. Neuron 109, 576–596.

Nones, J., Spohr, T. C. and Gomes, F. C. (2012). Effects of the flavonoid hesperidin in cerebral cortical progenitors in vitro: indirect action through astrocytes. Int J Dev Neurosci 30, 303–13.

Oberheim, N. A., Takano, T., Han, X., He, W., Lin, J. H., Wang, F., Xu, Q., Wyatt, J. D., Pilcher, W., Ojemann, J. G. et al. (2009). Uniquely hominid features of adult human astrocytes. J Neurosci 29, 3276-87.

Panatier, A., Vallee, J., Haber, M., Murai, K. K., Lacaille, J. C. and Robitaille, R. (2011). Astrocytes are endogenous regulators of basal transmission at central synapses. Cell 146, 785–98.

Park, J., Wetzel, I., Marriott, I., Dréau, D., D’Avanzo, C., Kim, D. Y., Tanzi, R. E. and Cho, H. (2018). A 3D human triculture system modeling neurodegeneration and neuroinflammation in Alzheimer’s disease. Nat Neurosci 21, 941–951.

Parslow, A., Cardona, A. and Bryson-Richardson, R. J. (2014). Sample drift correction following 4D confocal time-lapse imaging. J Vis Exp.

Perriot, S., Mathias, A., Perriard, G., Canales, M., Jonkmans, N., Merienne, N., Meunier, C., El Kassar, L., Perrier, A. L., Laplaud, D. A. et al. (2018). Human Induced Pluripotent Stem Cell-Derived Astrocytes Are Differentially Activated by Multiple Sclerosis-Associated Cytokines. Stem Cell Reports 11, 1199–1210.

Pestana, F., Edwards-Faret, G., Belgard, T. G., Martirosyan, A. and Holt, M. G. (2020). No Longer Underappreciated: The Emerging Concept of Astrocyte Heterogeneity in Neuroscience. In Brain Sciences, vol. 10.

Qiao, C., Yin, N., Gu, H. Y., Zhu, J. L., Ding, J. H., Lu, M. and Hu, G. (2016). Atp13a2 Deficiency Aggravates Astrocyte-Mediated Neuroinflammation via NLRP3 Inflammasome Activation. CNS Neurosci Ther 22, 451–60.

Quadri, M., Mandemakers, W., Grochowska, M. M., Masius, R., Geut, H., Fabrizio, E., Breedveld, G. J., Kuipers, D., Minneboo, M., Vergouw, L. J. M. et al. (2018). LRP10 genetic variants in familial Parkinson’s disease and dementia with Lewy bodies: a genome-wide linkage and sequencing study. Lancet Neurol 17, 597–608.

Reinhardt, P., Glatza, M., Hemmer, K., Tsytsyura, Y., Thiel, C. S., Hoing, S., Moritz, S., Parga, J. A., Wagner, L., Bruder, J. M. et al. (2013). Derivation and expansion using only small molecules of human neural progenitors for neurodegenerative disease modeling. PLoS One 8, e59252.

Rowitch, D. H. and Kriegstein, A. R. (2010). Developmental genetics of vertebrate glial-cell specification. Nature 468, 214–22.

Roybon, L., Lamas, N. J., Garcia, A. D., Yang, E. J., Sattler, R., Lewis, V. J., Kim, Y. A., Kachel, C. A., Rothstein, J. D., Przedborski, S. et al. (2013). Human stem cell-derived spinal cord astrocytes with defined mature or reactive phenotypes. Cell Rep 4, 1035–1048.

Santello, M., Toni, N. and Volterra, A. (2019). Astrocyte function from information processing to cognition and cognitive impairment. Nat Neurosci 22, 154–166.

Santos, R., Vadodaria, K. C., Jaeger, B. N., Mei, A., Lefcochilos-Fogelquist, S., Mendes, A. P. D., Erikson, G., Shokhirev, M., Randolph-Moore, L., Fredlender, C. et al. (2017). Differentiation of Inflammation-Responsive Astrocytes from Glial Progenitors Generated from Human Induced Pluripotent Stem Cells. Stem Cell Reports 8, 1757–1769.

Sardar, D., Cheng, Y.-T., Szewczyk, L. M., Deneen, B. and Molofsky, A. V. (2020). Chapter 32-Mechanisms of astrocyte development. In Patterning and Cell Type Specification in the Developing CNS and PNS (Second Editiong), (eds J. Rubenstein P. Rakic B. Chen and K. Y. Kwan), pp. 807–827: Academic Press.

Scheibye-Knudsen, M., Fang, E. F., Croteau, D. L., Wilson, D. M., 3rd and Bohr, V. A. (2015). Protecting the mitochondrial powerhouse. Trends Cell Biol 25, 158–70.

Schindelin, J., Arganda-Carreras, I., Frise, E., Kaynig, V., Longair, M., Pietzsch, T., Preibisch, S., Rueden, C., Saalfeld, S., Schmid, B. et al. (2012). Fiji: an open-source platform for biological-image analysis. Nat Methods 9, 676–82.

Schonhoff, A. M., Williams, G. P., Wallen, Z. D., Standaert, D. G. and Harms, A. S. (2020). Innate and adaptive immune responses in Parkinson’s disease. Prog Brain Res 252, 169–216.

Schwendy, M., Unger, R. E., Bonn, M. and Parekh, S. H. (2019). Automated cell segmentation in FIJI(R) using the DRAQ5 nuclear dye. BMC Bioinformatics 20, 39.

Siletti, K., Hodge, R., Mossi Albiach, A., Hu, L., Lee, K. W., Lönnerberg, P., Bakken, T., Ding, S.-L., Clark, M., Casper, T. et al. (2022). Transcriptomic diversity of cell types across the adult human brain. bioRxiv, 2022.10.12.511898.

Sloan, S. A., Darmanis, S., Huber, N., Khan, T. A., Birey, F., Caneda, C., Reimer, R., Quake, S. R., Barres, B. A. and Pasca, S. P. (2017). Human Astrocyte Maturation Captured in 3D Cerebral Cortical Spheroids Derived from Pluripotent Stem Cells. Neuron 95, 779–790 e6.

Sofroniew, M. V. and Vinters, H. V. (2010). Astrocytes: biology and pathology. Acta Neuropathol 119, 7–35.

Strokin, M., Seburn, K. L., Cox, G. A., Martens, K. A. and Reiser, G. (2012). Severe disturbance in the Ca2+ signaling in astrocytes from mouse models of human infantile neuroaxonal dystrophy with mutated Pla2g6. Hum Mol Genet 21, 2807–14.

Tchieu, J., Calder, E. L., Guttikonda, S. R., Gutzwiller, E. M., Aromolaran, K. A., Steinbeck, J. A., Goldstein, P. A. and Studer, L. (2019). NFIA is a gliogenic switch enabling rapid derivation of functional human astrocytes from pluripotent stem cells. Nat Biotechnol 37, 267–275.

Tcw, J., Wang, M., Pimenova, A. A., Bowles, K. R., Hartley, B. J., Lacin, E., Machlovi, S. I., Abdelaal, R., Karch, C. M., Phatnani, H. et al. (2017). An Efficient Platform for Astrocyte Differentiation from Human Induced Pluripotent Stem Cells. Stem Cell Reports 9, 600–614.

Torigoe, M., Yamauchi, K., Zhu, Y., Kobayashi, H. and Murakami, F. (2015). Association of astrocytes with neurons and astrocytes derived from distinct progenitor domains in the subpallium. Sci Rep 5, 12258.

Torres-Platas, S. G., Nagy, C., Wakid, M., Turecki, G. and Mechawar, N. (2016). Glial fibrillary acidic protein is differentially expressed across cortical and subcortical regions in healthy brains and downregulated in the thalamus and caudate nucleus of depressed suicides. Mol Psychiatry 21, 509–15.

Tsai, H. H., Li, H., Fuentealba, L. C., Molofsky, A. V., Taveira-Marques, R., Zhuang, H., Tenney, A., Murnen, A. T., Fancy, S. P., Merkle, F. et al. (2012). Regional astrocyte allocation regulates CNS synaptogenesis and repair. Science 337, 358–62.

van den Akker, E., Satchwell, T. J., Pellegrin, S., Daniels, G. and Toye, A. M. (2010). The majority of the in vitro erythroid expansion potential resides in CD34(-) cells, outweighing the contribution of CD34(+) cells and significantly increasing the erythroblast yield from peripheral blood samples. Haematologica 95, 1594–8.

Vanhauwaert, R., Kuenen, S., Masius, R., Bademosi, A., Manetsberger, J., Schoovaerts, N., Bounti, L., Gontcharenko, S., Swerts, J., Vilain, S. et al. (2017). The SAC1 domain in synaptojanin is required for autophagosome maturation at presynaptic terminals. Embo J 36, 1392–1411.

Verkhratsky, A. and Nedergaard, M. (2018). Physiology of Astroglia. Physiol Rev 98, 239–389.

Weissbein, U., Schachter, M., Egli, D. and Benvenisty, N. (2016). Analysis of chromosomal aberrations and recombination by allelic bias in RNA-Seq. Nat Commun 7, 12144.

Wu, T., Hu, E., Xu, S., Chen, M., Guo, P., Dai, Z., Feng, T., Zhou, L., Tang, W., Zhan, L. et al. (2021). clusterProfiler 4.0: A universal enrichment tool for interpreting omics data. The Innovation 2, 100141.

Xin, W., Schuebel, K. E., Jair, K. W., Cimbro, R., De Biase, L. M., Goldman, D. and Bonci, A. (2019). Ventral midbrain astrocytes display unique physiological features and sensitivity to dopamine D2 receptor signaling. Neuropsychopharmacology 44, 344–355.

Yang, Y., Song, J. J., Choi, Y. R., Kim, S. H., Seok, M. J., Wulansari, N., Darsono, W. H. W., Kwon, O. C., Chang, M. Y., Park, S. M. et al. (2022). Therapeutic functions of astrocytes to treat alpha-synuclein pathology in Parkinson’s disease. Proc Natl Acad Sci U S A 119, e2110746119.

Yun, S. P., Kam, T. I., Panicker, N., Kim, S., Oh, Y., Park, J. S., Kwon, S. H., Park, Y. J., Karuppagounder, S. S., Park, H. et al. (2018). Block of A1 astrocyte conversion by microglia is neuroprotective in models of Parkinson’s disease. Nat Med 24, 931–938.

Zaichick, S. V., McGrath, K. M. and Caraveo, G. (2017). The role of Ca(2+) signaling in Parkinson’s disease. Dis Model Mech 10, 519–535.

Zhou, Q., Viollet, C., Efthymiou, A., Khayrullina, G., Moritz, K. E., Wilkerson, M. D., Sukumar, G., Dalgard, C. L. and Doughty, M. L. (2019). Neuroinflammatory astrocytes generated from cord blood-derived human induced pluripotent stem cells. Journal of Neuroinflammation 16, 164.

