## Supplementary Material for "deCLUTTER^2+^ pipeline to analyze calcium traces in a novel stem cell model for ventral midbrain patterned astrocytes"

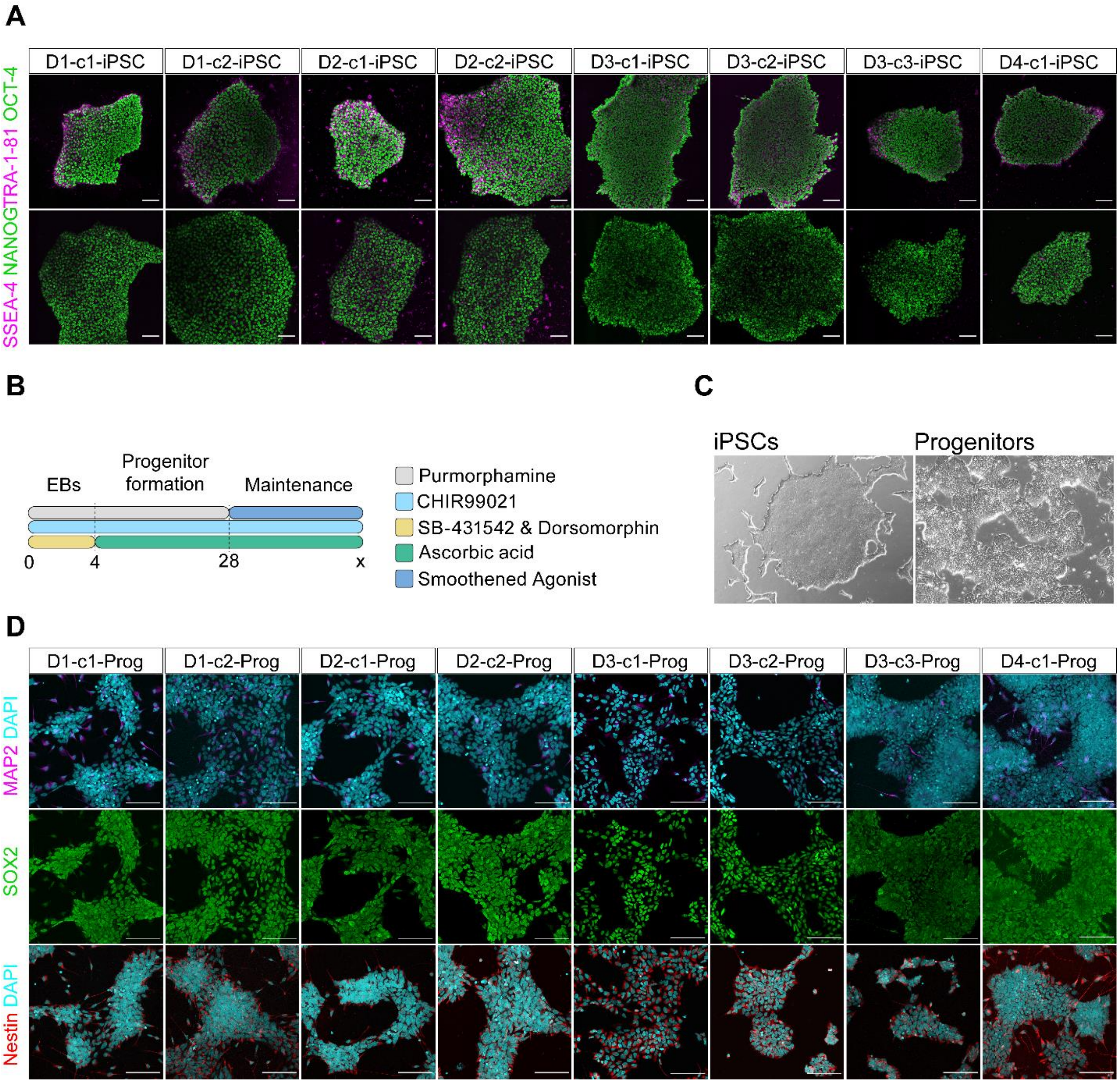

**Supplementary Fig. 1** Generation and characterization of iPSCs and iPSC-derived ventral midbrain patterned progenitors. **A.** Representative ICC images of eight iPSC lines staining positive for general pluripotency markers OCT4, TRA-1-81, NANOG, and SSEA-4. **B**. Schematic of the ventral midbrain progenitor differentiation protocol depicting the major steps with the accompanying supplements. **C**. Representative bright field images of feeder-free iPSC cultures and progenitor cultures. **D.** Representative ICC images of eight iPSC-derived ventral midbrain patterned progenitors staining positive for neural progenitor markers

SOX2 and Nestin and negative for mature neuron marker MAP2. Scale bars, 50 µm (in **C**) or 100 µm (in **D**).

Nuclei were counterstained with DAPI (cyan)

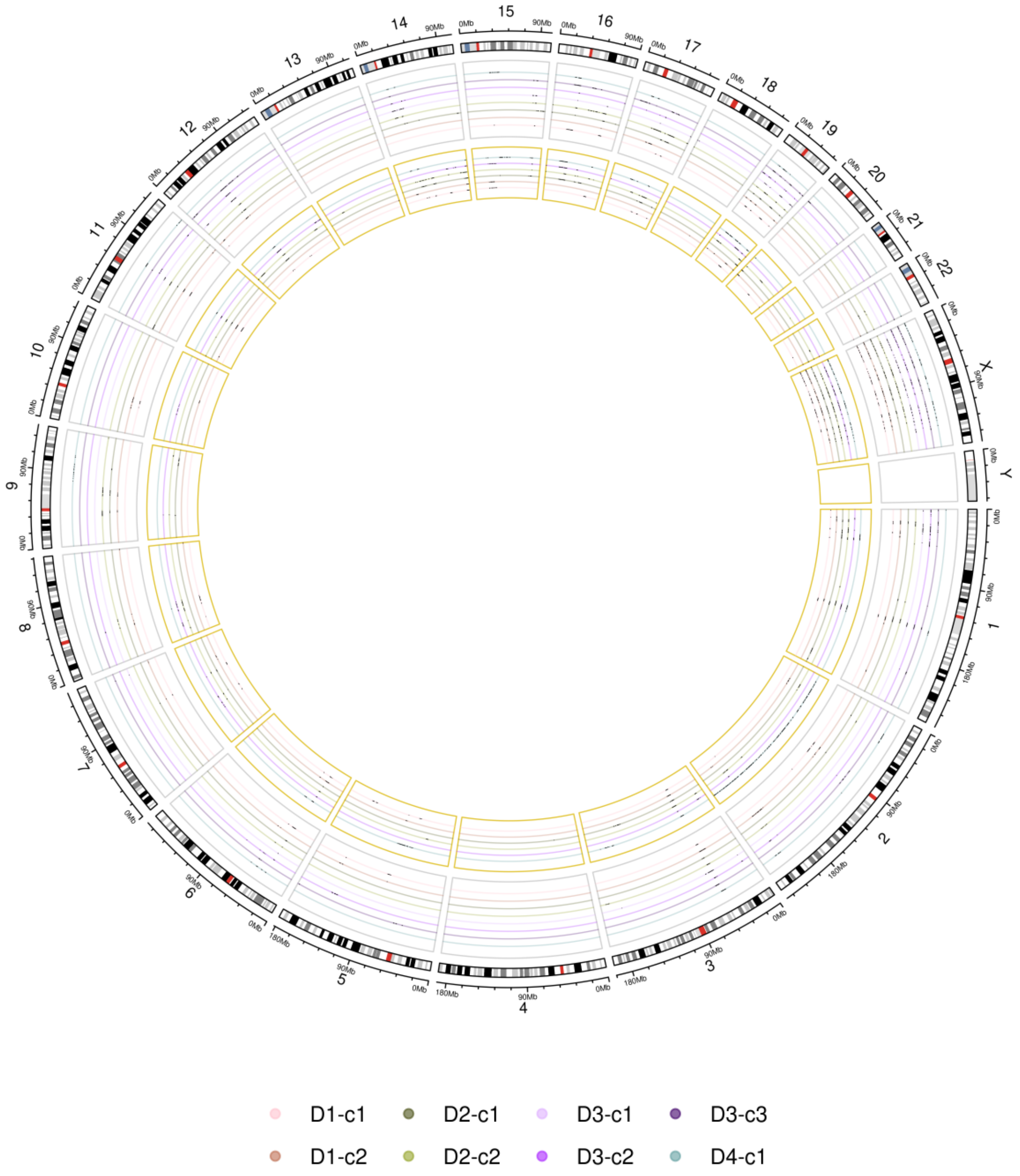

**Supplementary Fig. 2** eSNP-Karyotyping reveals chromosomal multiplication in one astrocyte line. Circle plot shows statistically significant multiplications detected by eSNP-Karyotyping (black dots). The grey inset reports the multiplications detected in the progenitors, while the golden inset those in the astrocytes.

Supplementary Table 1 Differentiation protocols to generate ventral midbrain astrocytes

| Reference | Model | Progenitors | Differentiation factors | Advantages | Potential limitations |
|---|---|---|---|---|---|
| | | Human iPSC-derived ventral midbrain astrocytes | | | |
| (Holmqvist et al., 2015) | 2D | Dual-SMAD inhibition and patterning toward a mesencephalic fate (floor-plate progenitors) via modulation of WNT and SHH signaling as described in (Kirkeby et al., 2012)<br><br>Supplements: LDN-193189, SAG, SB-431542, SHH-C25II N-terminus, CHIR-99021 | EGF, FGF-2, FBS | The high expression of astrocyte-specific markers: CD44, Connexin 43/GJA1, GLAST, NF-1A, and S100β; the bright GFAP expression can be maintained for several weeks after FACS; the secretion of pro-inflammatory cytokines and chemokines upon Il-1β and FBS treatment | The 130 days of culture; the necessity of the generation of the reporter line expressing TagRFP driven by the ABC1D element of the GFAP promoter (*$GFA_{ABC1D}$::TagRFP*) to obtain homogenous astrocyte populations through FACS |
| (Barbuti et al., 2020) | 2D | Dual-SMAD inhibition and patterning toward a mesencephalic fate (floor-plate progenitors) via modulation of WNT and SHH signaling, according to (Reinhardt et al., 2013)<br><br>Supplements: AA, CHIR-99021, dorsomorphin, PMA, SB-431542 | CNTF, EGF, FGF-2, FGF-8, LIF, Heparin, Heregulin, VPA | Serum-free; high expression of ALDH1L1, AQP4, Connexin 43/GJA1, GFAP, S100β, and Vimentin; spontaneous cytosolic $Ca^{2+}$ waves | The 120 days of culture; It requires many extrinsic gliogenic molecules |
| (de Rus Jacquet, 2019) (de Rus Jacquet et al., 2021) | 2D | Dual-SMAD inhibition and patterning toward a mesencephalic fate (floor-plate progenitors) via modulation of WNT and SHH signaling as described in (Kriks et al., 2011)<br><br>Supplements: CHIR-99021, LDN193189, Purmorphamine, SB-431542, SHH-C25II N-Terminus | Commercial astrocyte medium (ScienCell) | Astrocytes are obtained in a relatively short period (~28 days) | The medium contains 2% FBS; the cells have a flat morphology; the poor immunocytochemical and functional characterization of the obtained astrocytes |
| (Crompton et al., 2021) | 2D | Dual-SMAD inhibition and patterning toward a mesencephalic fate (floor-plate progenitors) via modulation of WNT and SHH signaling | BMP4, EGF, LIF | High expression of GFAP, S100β, and ventral midbrain identity markers FOXA2, LMX1B; | Time-consuming (120+ days); limited functional characterization |

| | | |
|---|---|---|
| | Supplements: LDN193189, SB-431542, CHIR-99021, SHH-C25II N-Terminus | morphological changes and a significant increase in the secretion of IL-6 upon treatment with pro-inflammatory cytokines IL-1α or IL-1β |

NF-1A, nuclear factor-1; FACS, fluorescence-activated cell sorting; RFP, red fluorescence protein; PMA, purmorphamine; AA, ascorbic acid; FGF-8, fibroblast growth factor 8; VPA, valproic acid; TGF-β, transforming growth factor beta; cAMP, adenosine-3′,5′-cyclic monophosphate; dbcAMP, dibutyryl-cAMP; SAG, smoothened agonist; NT-3, neurotrophin-3; DHA, docosahexaenoic acid

Supplementary Table 2 Cell culture media composition

| **Astrocyte basal media (ABM).** Keep up to one month at 4°C. Do not freeze and thaw! | | |
|---|---|---|
| DMEM-F12, GlutaMAX | 100% (500 mL) | Gibco™, 10565018 |
| N2 (100×) | 1× (5 mL) | Gibco™, 17502001 |
| B27 without vitamin A (50×) | 1× (10 mL) | Gibco™, 12587010 |
| PenStrep (100×) | 1× (5 mL) | Gibco™, 15070063 |
| MEM non-essential amino acids (NEAA) solution (100×) | 1× (5 mL) | Gibco™, 11140050 |

| **Glial expansion media** = ABM + supplements; added at the time of media change | | |
|---|---|---|
| HEPES | 10 mM | Gibco™, 15630056 |
| EGF | 10 ng/mL | Peprotech, AF-100-15 |
| FGF-2 (bFGF) | 10 ng/mL | Peprotech, 100-18B |

| **Glial induction media** = ABM + supplements; added at the time of media change | | |
|---|---|---|
| HEPES | 10 mM | Gibco™, 15630056 |
| EGF | 10 ng/mL | Peprotech, AF-100-15 |
| LIF | 10 ng/mL | Peprotech, 300-05 |

| **Glial maturation media** = ABM + supplements; added at the time of media change | | |
|---|---|---|
| HEPES | 10 mM | Gibco™, 15630056 |
| CNTF | 10 ng/mL | Peprotech, 450-13 |

| **Glial maintenance media** = ABM + supplements; added at the time of media change | | |
|---|---|---|
| HEPES | 10 mM | Gibco™, 15630056 |

Supplementary Table 3 Details of the primary antibodies used in this study

| Epitope | Catalog | Source | Host | Dilution ICC |
|---|---|---|---|---|
| AQP-4 | HPA014784 | Atlas Antibodies | Rabbit | 1:200 |
| CD44 (F10-44-2) | AB6124 | Abcam | Mouse | 1:200 |
| GFAP | N206A/8 | DSHB | Mouse | 1:200 |
| GFAP | AB5804 | Millipore Sigma | Rabbit | 1:200 |
| NANOG | AB21624 | Abcam | Rabbit | 1:75 |
| Nestin (10C2) | MAB5326 | Millipore Sigma | Mouse | 1:200 |
| MAP2 | 188 004 | Synaptic Systems | Guinea pig | 1:250 |
| OCT4 | AB19857 | Abcam | Rabbit | 1:250 |
| S100β | HPA015768 | Sigma-Aldrich | Mouse | 1:100 |
| SOX2 | AB5603 | Millipore Sigma | Rabbit | 1:200 |
| SSEA4 | AB16287 | Abcam | Mouse | 1:75 |
| TRA1-81 | AB16289 | Abcam | Mouse | 1:75 |

Supplementary Table 4 Details of the 14 stem cell-derived clonal lines used in this study

| Donor ID | Source | Donor Age | Donor Sex | Donor Sampling Site | Progenitor Clone IDs | Astrocyte Clone IDs | RIN for all the clones |
|---|---|---|---|---|---|---|---|
| **D1** | Gibco™, Lot number: 1903939 | 33 | Female | Dermal Fibroblast | c1, c2 | c1, c2 | 10 |
| **D2** | Gibco™, Lot number: 181388 | 34 | Female | Dermal Fibroblast | c1, c2 | c1, c2 | 10 (only D2-c2-Astro RIN= 8.2) |
| **D3** | *in house* | 68 | Female | Dermal Fibroblast | c1, c2, c3 | c3 | 10 |
| **D4** | *in house* | 83 | Female | Erythroid Progenitors | c1 | c1 | 10 |

Supplementary Table 5 qPCR primer sequences

| Primer name | Sequence |
|---|---|
| Reactivity-specific genes | |
| *C3*-FW | AAAAGGGGCGCAACAAGTTC |
| *C3*-RV | GATGCCTTCCGGGTTCTCAA |
| *GFAP*-FW | AGAAGCTCCAGGATGAAACC |
| *GFAP*-RV | AGCGACTCAATCTTCCTCTC |
| *LCN2*-FW | ATCACCCTCTACGGGAGAACC |
| *LCN2*-RV | ACTCAGCCGTCGATACACTG |
| *SERPINA3*-FW | TGCCAGCGCACTCTTCATC |
| *SERPINA3*-RV | TGTCGTTCAGGTTATAGTCCCTC |
| Housekeeping genes | |
| *CLK2*-FW | TCGTTAGCACCTTAGGAGAGG |
| *CLK2*-RV | TGATCTTCAGGGCAACTCG |
| *COPS5*-FW | CCAGGAACCATTTGTAGCAG |
| *COPS5*-RV | GTAGCCCTTTGGGTATGTCC |
| *RNF10*-FW | GCATCTGTGAACTGGCTTTG |
| *RNF10*-RV | CTGACGTTTCCTCTTCTCAATG |

### References

**Barbuti, P. A., Antony, P., Novak, G., Larsen, S. B., Berenguer-Escuder, C., Santos, B. F. R., Massart, F., Grossmann, D., Shiga, T., Ishikawa, K.-i. et al.** (2020). IPSC-derived midbrain astrocytes from Parkinson’s disease patients carrying pathogenic <em>SNCA</em> mutations exhibit alpha-synuclein aggregation, mitochondrial fragmentation and excess calcium release. *bioRxiv*, 2020.04.27.053470.

**Crompton, L. A., McComish, S. F., Stathakos, P., Cordero-Llana, O., Lane, J. D. and Caldwell, M. A.** (2021). Efficient and Scalable Generation of Human Ventral Midbrain Astrocytes from Human-Induced Pluripotent Stem Cells. *J Vis Exp*.

**de Rus Jacquet, A.** (2019). Preparation and Co-Culture of iPSC-Derived Dopaminergic Neurons and Astrocytes. *Curr Protoc Cell Biol* **85**, e98.

**de Rus Jacquet, A., Tancredi, J. L., Lemire, A. L., DeSantis, M. C., Li, W. P. and O'Shea, E. K.** (2021). The LRRK2 G2019S mutation alters astrocyte-to-neuron communication via extracellular vesicles and induces neuron atrophy in a human iPSC-derived model of Parkinson's disease. *Elife* **10**.

**Holmqvist, S., Brouwer, M., Djelloul, M., Diaz, A. G., Devine, M. J., Hammarberg, A., Fog, K., Kunath, T. and Roybon, L.** (2015). Generation of human pluripotent stem cell reporter lines for the isolation of and reporting on astrocytes generated from ventral midbrain and ventral spinal cord neural progenitors. *Stem Cell Res* **15**, 203-20.

**Kirkeby, A., Grealish, S., Wolf, D. A., Nelander, J., Wood, J., Lundblad, M., Lindvall, O. and Parmar, M.** (2012). Generation of regionally specified neural progenitors and functional neurons from human embryonic stem cells under defined conditions. *Cell Rep* **1**, 703-14.

**Kriks, S., Shim, J. W., Piao, J., Ganat, Y. M., Wakeman, D. R., Xie, Z., Carrillo-Reid, L., Auyeung, G., Antonacci, C., Buch, A. et al.** (2011). Dopamine neurons derived from human ES cells efficiently engraft in animal models of Parkinson's disease. *Nature* **480**, 547-51.

**Reinhardt, P., Glatza, M., Hemmer, K., Tsytsyura, Y., Thiel, C. S., Hoing, S., Moritz, S., Parga, J. A., Wagner, L., Bruder, J. M. et al.** (2013). Derivation and expansion using only small molecules of human neural progenitors for neurodegenerative disease modeling. *PLoS One* **8**, e59252.
